# *Neisseria leonis* sp. nov. isolated from rabbits, reclassification of *Uruburuella suis, Uruburuella testudinis, Kingella potus, Bergeriella denitrificans* and *Morococcus cerebrosus* into *Neisseria* genus and reclassification of *Neisseria shayeganii* into *Eikenella* genus

**DOI:** 10.1101/2023.02.24.529859

**Authors:** M. Boutroux, S. Favre-Rochex, O. Gorgette, G. Touak, E. Muhle, O. Chesneau, D. Clermont, P. Rahi

## Abstract

Genome sequence-based identification of two strains (3986^T^ and 51.81) isolated from rabbits in France in 1972 and 1981 and deposited in the Collection of Institut Pasteur (CIP) has led to the description of a novel species in the genus *Neisseria*. The cells of both strains were non-motile, Gram-stain-negative and diplococcobacilli. Optimal growth on trypticase soy agar was recorded at 37°C and pH 8.5 in aerobic conditions. Phylogeny based on 16S rRNA gene placed the strains close to *Neisseria bacilliformis* ATCC BAA-1200^T^ (96.38%) nesting with the members of Neisseriaceae family. Furthermore, the phylogenetic analysis based on *bac120* gene set from the Genome Taxonomy Database (GTDB) placed both strains within the monophyletic *Neisseria* clade, which also included type strains of *Morococcus cerebrosus, Bergeriella denitrificans*, *Kingella potus, Uruburuella suis* and *Uruburuella testudinis*. However, *Neisseria shayeganii* strain 871^T^ was placed outside *Neisseria* clade and close to the members of *Eikenella* genus. Strains 3986^T^ and 51.81 were placed in a branch distinct from all species of the genus *Neisseria* and exhibited the average nucleotide identity (ANI) and digital DNA–DNA hybridization (dDDH) values below the species demarcation values. In contrast, ANI value within the two strains was 96.9% confirming that they represent same species. The genomic DNA G+C content of strain 3986^T^ was 56.92%. Based on the phylogenetic and phenotypic data, the strains 3986^T^ and 51.81 represent a novel species of the genus *Neisseria*, for which the name *Neisseria leonis* sp. nov. is proposed (type strain 3986^T^ = CIP 109994^T^ = LMG 32907^T^). Additionally, based on phylogenetic analysis, DUS dialect and average amino acid identity (AAI) values, we also proposed the reclassification of *Morococcus cerebrosus*, *Bergeriella denitrificans*, *Kingella potus, Uruburuella suis* and *Uruburuella testudinis* into *Neisseria* genus and *Neisseria shayeganii* into *Eikenella* genus.

**Author Notes:** The GenBank accession numbers for the 16S rRNA gene sequence of strains 3986^T^ and 51.81 are respectively OQ121838.1 and OQ428162.1. The draft genome sequences have been deposited in GenBank under the accession numbers JAPQFK000000000 (strain 3986^T^) and JAPQFL000000000 (strain 51.81).

Further explanations mentioned in the article as well as 7 supplementary tables and 7 supplementary figures are available with the online version of this article.

## Introduction

The genus *Neisseria* comprises 29 validly published species along with the important human pathogens *Neisseria gonorrhoeae* and *Neisseria meningitidis* according to the List of Prokaryotic names with Standing in Nomenclature (LPSN) [1]. Most *Neisseria* species have been isolated from a wide range of animals like Arctican greater white-fronted goose, Californian sea lion, and Tibetan plateau pika [2–4], while a few are common in oral and nasopharyngeal microbiotes in humans and can even be opportunistic pathogens [5]. *Neisseria* species isolated from herbivorous mammals includes *Neisseria animalis* from the pharyngeal mucosa of the guinea pig [6], *Neisseria dentiae* from the dental plaque of a dairy cow [7] and most recently *Neisseria musculi* from the oral cavity of the wild house mouse [8].

The strains 3986^T^ and 51.81 were isolated from rabbits at Institut Pasteur Lyon and Tours Departmental Veterinary Laboratory and deposited at the Collection of Institut Pasteur (CIP) in 2002 and in 1987, respectively. Prior to genome sequence-based identification, both were assigned as *Flavobacterium* sp. Culture collections are the reliable source of providing quality controlled microbial resources, which are often used by researchers as reference material or as model system to understand complex biological questions. It is expected that these two strains will be used in several studies as reference cultures to investigate antibiotic resistance and virulence factors in *Neisseria* to understand the ecology and evolution of this group. Additionally, rabbits are often used as animal models, and we anticipate that strains 3986^T^ and 51.81 could be used to understand the several aspects of host-microbe interactions where rabbits are used as animal models. *Neisseria musculi* isolated from mice has already been used extensively for studying commensal colonization [9,10]. Based on detailed genomic studies and polyphasic characterization, we purposed that strain 3986^T^ and 51.81 represent a new *Neisseria* species. Our study, therefore, contributes to improve the understanding of the biodiversity of the genus *Neisseria*.

## Materials and methods

### Isolation

Strain 3986^T^ (= R726^T^ = CIP 109994^T^) was isolated at the Institut Pasteur Lyon in 1972 from the liver of a baby rabbit. Strain 51.81 (= CIP 103045) was isolated by the Tours Departmental Veterinary Laboratory in 1981 from the lung of a rabbit. Both strains were revived from the lyophilized vials on Trypticase Soy Agar (TSA) at 30°C for 24 hours. Strains were stored both as lyophilized vials with 5% myo-inositol and 10% skimmed milk at 4°C and with 15% glycerol at −80°C.

### 16S rRNA gene phylogeny

For 16S rRNA gene sequencing, DNA was extracted from the strains using the InstaGenMatrix (Bio-Rad, USA) and stored at −20°C. The 16S rRNA gene was amplified using primers 27F-YM 5’-AGAGTTTGATYMTGGCTCAG-3’ and 1391R 5’-GACGGGCGGTGWGTRCA-3’ and GoTaq DNA Polymerase (Promega, USA). The PCR was performed as follows: initial denaturation at 95°C for 3 minutes and then 30 cycles of denaturation (95°C, 45s), hybridization (60°C, 45s), elongation (72°C, 90s). The amplified product was sent for Sanger sequencing at Eurofins Genomics (Germany). The sequence quality was checked with BioEdit 7.2.5, and trimming and assembly were performed using CLCGenomicsWorkbench 20.0.4.

The sequences were searched against the NCBI nucleotide database using BLASTn search tool to find the sequences of closely related strains. Additionally, rabbit microbiome data from a 2019 study [11] was also searched to find Amplicon Sequence Variants (ASVs) identical to strain 3986^T^ (see supplementary text 1 for detailed procedure).

16S rRNA sequences of strains 3986^T^ and 51.81 were run against EZBioCloud database using 16S-based ID tool [12]. Sequences of 16S rRNA gene from type strains of the closely related species resulted from EZBioCloud search and from the additional strains resulted from NCBI database search were used to construct phylogenetic trees. All sequences were aligned using MUSCLE [13] in MEGA 11 [14], and phylogenetic trees were built using maximum likelihood (ML), neighbor joining (NJ) and maximum parsimony (MP), with 1000 bootstrap replications.

### Genome sequencing and phylogenetic analysis

For genome sequencing, DNA was extracted using the Wizard Genomic DNA purification kit (Promega, USA) and stored at −20 °C. The sequencing was performed using a Nextseq 500 Instrument (Illumina, USA) with a 2×150 bp paired-end protocol at the Plateforme de Microbiologie Mutualisée (P2M) of the Institut Pasteur. The reads were assembled using the pipeline fq2dna 21.06 (gitlab.pasteur.fr/GIPhy/fq2dna). Genomic characteristics and quality were evaluated with contig_info 2.1 [15] and checkM 1.1.3 [16].

A dataset of 79 genomes consisting of the genomes of strains 3986^T^ and 51.81, type strains of *Neisseria* species, type strains of species from other genera of the Neisseriaceae family close to *Neisseria* genus and several unassigned species closely related to *Neisseria* mentioned in GTDB [17] (see supplementary text 2 for detailed procedure). We used GTDBTk 2.1.1 [18] to identify, extract and align *bac120* marker gene set from all the genomes. The resulted alignment was used to build a maximum likelihood phylogenetic tree using IQ-TREE 2.2.2.2 [19]. Another phylogenetic analysis was performed using the aligned sequences of 53 genes, identified and extracted by using the rMLST scheme [20] of BIGSdb [21] and aligned with MAFFT [22]. The construction of phylogenetic trees involved the selection of the best method by ModelFinder [23] and 1000 ultrafast bootstrap replications. Average nucleotide identities (ANIs) between the genomes of the dataset were calculated with FastANI [24]. The dDDH values were computed with TYGS [25]. The count function of the Jellyfish 2.3 module [26] was used to count the number of repeats of each k-mer of size 12 and evaluate the DNA Uptake Sequence (DUS) dialects used by the genomes of the dataset.

The EzAAI pipeline [27] was used to calculate AAI values and a heatmap was created with the results with the heatmap.2 function of the gplots module in R. Strains 3986^T^ and 51.81 were submitted to VFanalyzer [28] to identify virulence factors and resistance genes search was done on CARD with PGAP [29] annotated files using the Resistance Gene Identifier (RGI) tool [30]. The biosynthetic gene clusters (BGCs) for various secondary metabolites were identified by using an online genome mining pipeline antiSMASH 7.0 [31].

### Physiology

The International Code of Nomenclature of Prokaryotes (ICNP) subcommittee for the family *Neissericeae* has been inactive since 1982 and there are no minimal standards. Therefore, we performed the phenotypic characteristics on the basis of recent *Neisseria* species descriptions [3,4,8,32]. Strains were grown on different growth media, including TSA plates, Columbia agar with 10% horse blood and Brain Heart Infusion (BHI). Temperature range was determined using TSA plates incubated at different culture conditions (4, 15, 28, 37 and 45°C). All later incubations were done at 37°C unless mentioned otherwise. Tryptone soya broth (TSB) with different pH (4, 5.5, 7, 8.5, 10, 11 and 12) set by adding HCl and NaOH before autoclaving was used to estimate the growth in different pH. Similarly, TSB pH 7 with different concentrations of oxgall (0.1%, 0.3% and 0.5%) was used to determine the bile salt tolerance. Growth was measured by measuring absorbance at 600 nm with the Infinite M Nano+ (TECAN, Switzerland). Incubation under microaerophilic and anaerobic conditions were tested using round jars of 2.5L with CampyGen and AnaeroGen products (ThermoFisher, USA).

Microscopic observations were performed to determine the cell shape, size and motility of cultures grown on BHI medium. Motility was determined examining a small drop of bacterial culture in center of a microscope slide with the wet mount method [7]. Gram-staining was performed by using Aerospray Gram (ELITechGroup, France). For electronic microscopy, the bacterial cells were fixed to 300-mesh Formvar-Cu-coated grids (Electron Microscopy Sciences, UK) for 15 minutes. The grids with cells were placed on a drop of ultrapure water and transferred to a drop of 2% glutaraldehyde in 0.1M sodium cacodylate buffer for 10 minutes at room temperature. After rinsing with ultrapure water, grids were stained for 15 sec with 2% aqueous uranyl acetate and dried. Images were captured with Tecnai Spirit 120Kv transmission electron microscope (TEM) equipped with a bottom-mounted Eagle 4kx4k camera (FEI, USA).

Cultures for biochemical tests were grown using TSA plates incubated for 24 hours unless specified otherwise. Oxidase activity was determined by putting a smear of culture on an oxidase strip (Sigma-Aldrich, USA), and catalase activity was done recording the bubble formation by bacterial cells, in response to hydrogen peroxide solution (Sigma-Aldrich, USA).

Reference strains chosen for additional tests were the closest species to strains 3986^T^ and 51.81 according to ANI and TYGS (*Neisseria dentiae* CIP 106968^T^) as well as the 3 species clustering with strains 3986^T^ and 51.81 in the *bac120* tree (*Kingella potus* CIP 108935^T^, *Neisseria bacilliformis* DSM 23338^T^ and *Neisseria elongata* CIP 72.27^T^.) API NH and API ZYM (bioMérieux, France) were done with *Neisseria gonorrhoeae* type strain CIP 79.18^T^ in addition to the other strains. Strains were grown on chocolate agar for the API NH following manufacturer’s instructions and also for API ZYM in the case of *Neisseria gonorrhoeae* CIP 79.18^T^. In addition, *Neisseria bacilliformis* DSM 23338^T^ had to be grown at 37°C in aerobic conditions for all experiments. Nitrate and nitrite reduction tests were done incubating the bacteria in nitrate and nitrite broth before adding the reagents NIT 1 and NIT 2 (bioMérieux, France) as well as zinc for the nitrate reduction test. The antibiotic susceptibility was determined using the disc diffusion method on Mueller–Hinton agar for 16 antibiotics: Amoxicillin 20 μg, Cefotaxin 5 μg, Chloramphenicol 30 μg, Ciprofloxacin 5 μg, Eythromycin 15 μg, Fosfomycin 200 μg, Fusidic Acid 10 μg, Lincomycin 15 μg, Mecillinam 10 μg, Penicillin G 6 μg, Rifampicin 5 μg, Streptomycin 10 μg, Sulfonamides 200 μg, Tetracycline 30 μg, Trimethoprim 25 μg, and Vancomycin 5 μg.

## Results

### 16S rRNA analysis and phylogeny

The 16S rRNA gene of strains 3986^T^ and 51.81 sequenced by Sanger method were 1254 bp and 1263 bp long, while 1536 bp long sequences were extracted from the genome sequences of both strains. The sequences originated from different sequencing methods shared 100% identity. Therefore, we used the 16S rRNA gene sequences extracted from genomes for all database searches and phylogenetic analysis. The 16S rRNA gene sequences of both strains exhibited 99.09% identity when compared to each other, which is above the threshold for species delineation of 98.65% [33]. Additionally, the search in NCBI nucleotide database resulted 99.05% identity with the 16S rRNA gene sequence of the *Neisseria* sp. strain CCUG 45853 (AY064548.1), indicating its close similarity to the strain 3986^T^ and they are member of same species. Interestingly, the strain CCUG 45853 was isolated from the nose of a rabbit with symptoms of cold in Berne, Switzerland (http://www.ccug.se/strain?id=45853).

This intrigued us to further investigate the proximity between this species and rabbits, and we searched for identical ASVs in the 16S rRNA gene amplicon sequences data from rabbits and other animals [11,34]. During this, we realize the difficulties in accessing the microbiome sequence projects, particularly microbiome of different organs of rabbit [34], for which an accession number (PRJCA006344) was provided in the study, but did not exist in the databases and authors never replied to our requests. However, we used the gut microbiome data from Spanish rabbits to find identical 16S rRNA gene amplicons (SRP129755) [11]. We could find one ASV which was 100% identical to strain 3986^T^ and was present 19 times in one of the 126 samples (SRR6475038) (the metadata concerning this ASV is available in the supplementary material). The strains 3986^T^ and 51.81 were isolated from the liver and the lung of rabbits, and their lower occurrence in the fecal microbiome suggested the possible specificity of these strains to colonize vital organs of rabbits. It must be mentioned that the 16S rRNA gene V4 region (253 nucleotides) of *Kingella potus* and strain 3986^T^ are 100% identical, so the conclusions based on such microbiome analysis can be misleading.

EZBioCloud returned 37 hits (Table S1) above the 16S threshold for genus delineation of 94.5% [35]. The closest hits included type strains of 26 species of *Neisseria* and the members of other genera like *Kingella, Bergeriella, Morococcus, Eikenella, Conchiformibius* and *Alysiella*. The closest hit with strain 3986^T^ was *Neisseria bacilliformis* strain ATCC BAA-1200^T^ with 96.38% identity. All trees placed strains 3986^T^, 51.81 and CCUG 45853 in one distinct clade. In the neighbor-joining tree they were clustered with *Neisseria bacilliformis* strain ATCC BAA-1200^T^ (Fig. 1), but the bootstrap value was below 50% and this phylogenetic placement was not consistent with the trees obtained with the maximum likelihood and the maximum parsimony methods (Figs S1 and S2). Additionally, in the 16S rRNA gene phylogeny, the members of *Neisseria* do not form a monophyletic group, as the members of *Kingella, Bergeriella* and *Morococcus* are often nested within the group of *Neisseria* (Fig. 1).

**Fig. 1:**
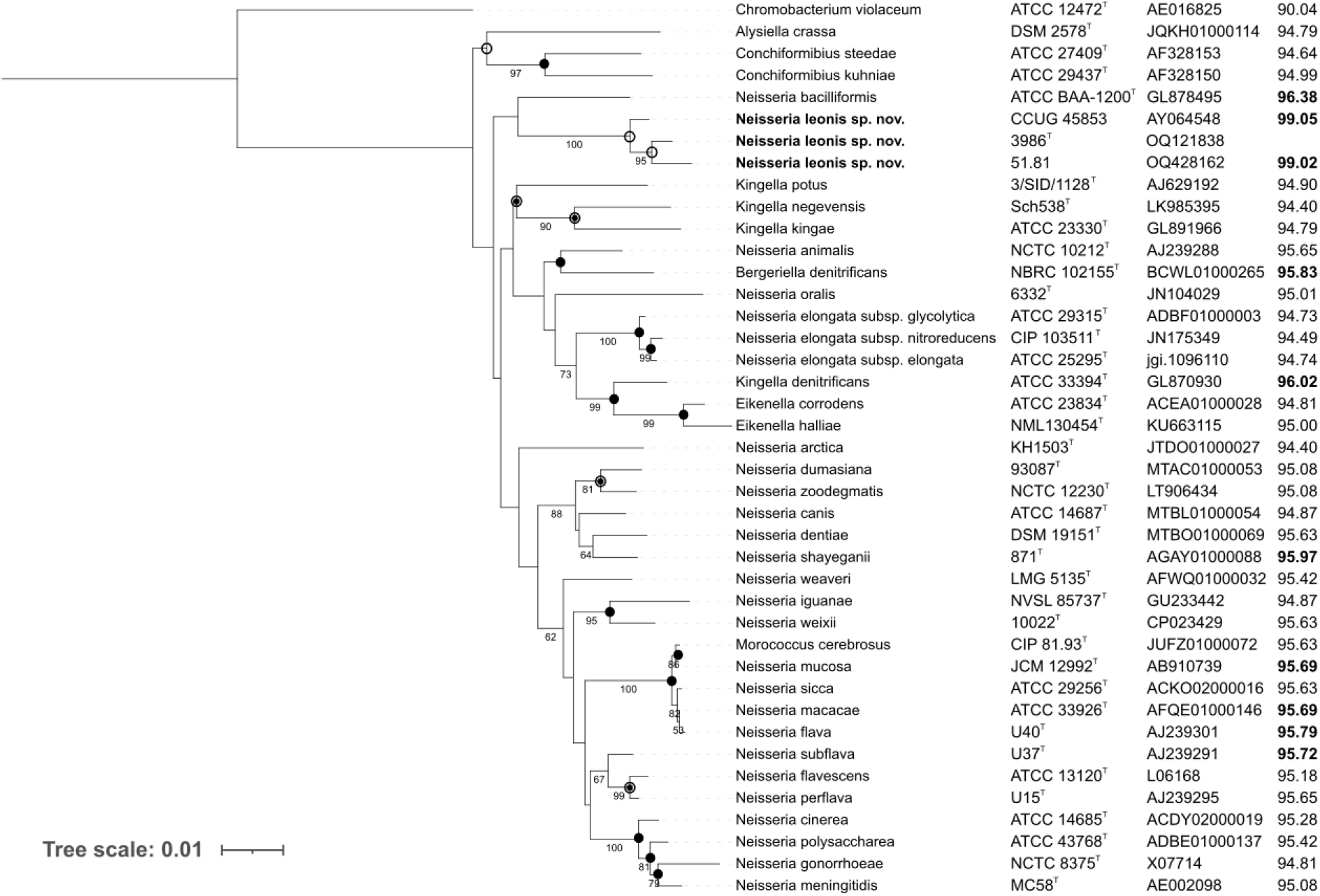
16S rRNA gene phylogenetic tree obtained in MEGA 11 with neighbor-joining method from the sequences of strains 3986^T^, 51.81, CCUG 45853 and 37 top hits obtained from EZBioCloud with strain 3986^T^. Last column shows % of identity with the 16S rRNA gene sequence of strain 3986^T^ *Chromobacterium violaceum* ATCC 12472^T^ was used as an outgroup. Bootstrap values above 50% are displayed. Empty circles indicate branches that were also found using the maximum likelihood method, circles with a dot indicate branches that were also found using the maximum parsimony method and filled circles indicate branches that were also found using both methods. Bar, 0.01 changes per site.

### Genome features

The assembly of the genome of strain 3986^T^ resulted 22 contigs, with a N50 of 137808 bp and a 62.6X coverage, while strain 51.81 had 29 contigs, with a N50 of 471292 bp and a 61.1X coverage. DNA G+C content of strains 3986^T^ and 51.81 was 56.92 and 59.96% respectively and was at the upper fringe of the G+C content of *Neisseria* (44.6% to 59.4%). Only three *Neisseria* species had a higher G+C content. Sequence length of strains 3986^T^ and 51.81 was 2.18 Mb and 2.29 Mb, which is in accordance with the lengths of the 59 *Neisseria* genomes (average: 2.45 Mb, standard deviation: 0.2 Mb). Strains 3986^T^ and 51.81 shared an ANI of 96.96% (reversed identity of 96.91%) and a dDDH of 70.8%, which were slightly above the thresholds for species delineation of 95-96% for ANI and 70% for dDDH [35]. ReferenceSeeker and TYGS against those two genomes returned similar results compared to EZBioCloud, with additional hits from *Uruburuella* genus (Table S2). The closest hit for strain 3986^T^ was *Neisseria dentiae* DSM 19151^T^ for ANI and dDDH values with 79.89% and 23.8% identities, respectively.

The core-gene tree based on *bac120* gene (Fig. 2) provided genus-specific grouping, while rMLST tree (Fig. S3) was unable to separate the different genera into distinctive clusters. More specifically, *Alysiella, Simonsiella, Kingella, Conchiformibius* and *Eikenella* species were placed within the main *Neisseria* cluster. In the past, rMLST was described as the best method for classifying *Neisseria* [36]. However, we found that *bac120* marker set which also includes most of the genes in the rMLST scheme and several other genes is more discriminatory. The strains 3986^T^ and 51.81 were always clustered together and were placed within the *Neisseria* group (Fig. 2). In addition to this, the placement of five non-*Neisseria* species including *Morococcus cerebrosus*, *Bergeriella denitrificans*, *Kingella potus, Uruburuella suis* and *Uruburuella testudinis* within *Neisseria* and placement of *Neisseria shayeganii* within *Eikenella* exhibited the possibility of reclassification of these taxa (see Fig. S4 for the ANI values between the type strains of those species and the other genomes of the dataset). The rMLST tree also corroborated these observations (Fig. S3).

**Fig. 2:**
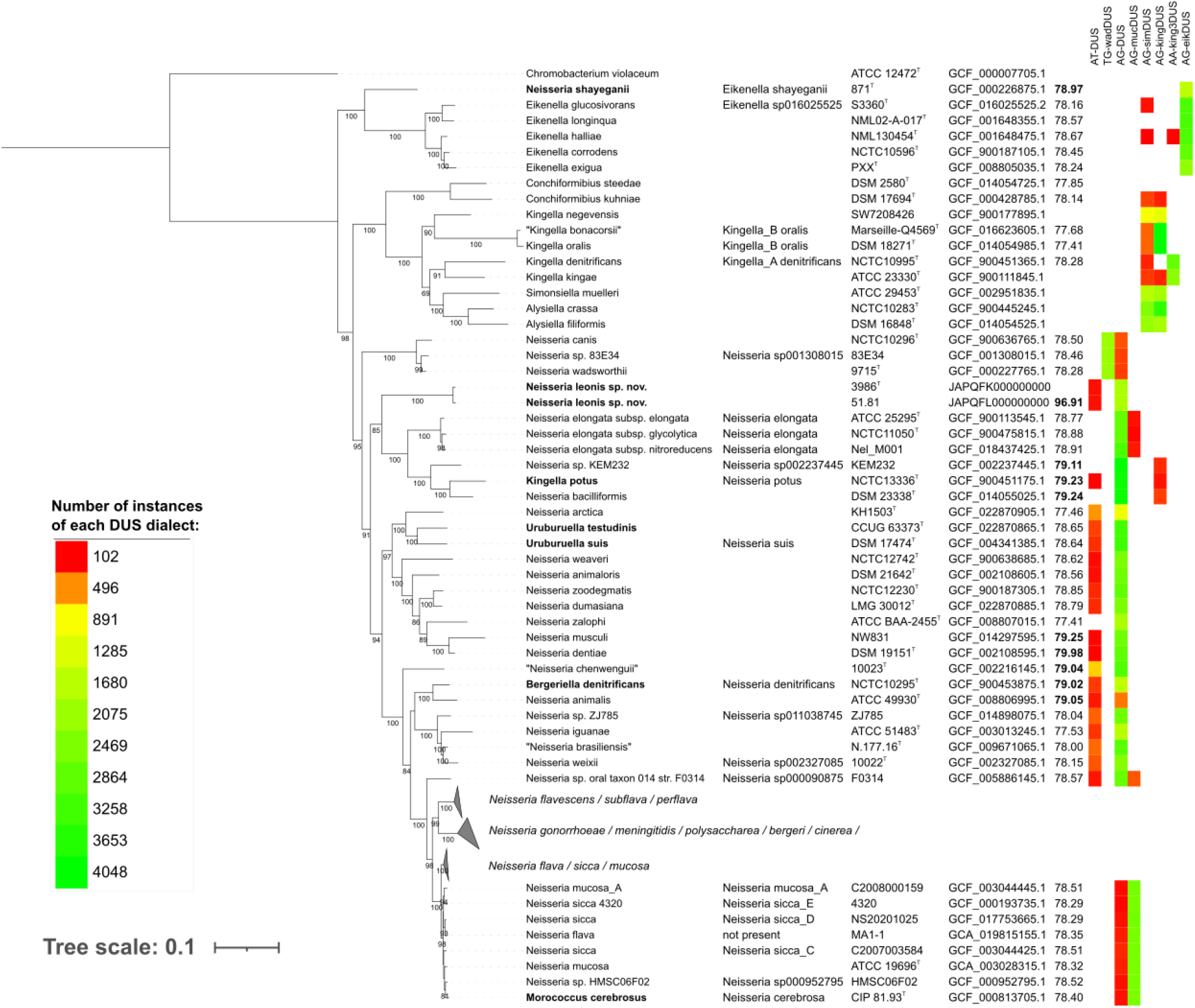
Phylogenetic tree obtained from the alignment of the *bac120* genes with maximum likelihood method. Each line contains the name of the species according to LPSN and according to GTDB if it differs, the isolate name, the accession number, the highest ANI values with strain 3986^T^ (top 10 values are in bold) and a heatmap depicting the number of instances of each DUS dialect (for numbers>100). Three *Neisseria* clades are collapsed here because they are not relevant for the species that concerned us. *Chromobacterium violaceum* ATCC 12472^T^ was used as an outgroup. Bootstrap values above 50% are displayed. Bar, 0.1 changes per site.

A detailed analysis of DUS dialects further supported these reclassifications (Fig. 2). The dominance of one variant or another is consistent with a phylogeny of Neisseriaceae species based on core genome sequences [37]. The DUS dialects used by each species confirmed the accuracy of the clusters in the *bac120* tree. Strains 3986^T^, 51.81, type strains of *Uruburuella* species, *Kingella potus* and *Bergeriella denitrificans* used the AG-DUS dialect, a dialect typical of *Neisseria*. Type strain of *Morococcus cerebrosus* used another *Neisseria* dialect, AG-mucDUS, while type strain of *Neisseria shayeganii* employed AG-eikDUS, the DUS dialect specific of *Eikenella* species. The detail of the DUS sequence counts is available in the supplementary material (Table S4).

Amino Acid Identity (AAI) values computed for 20 genomes including the strains 3986^T^, 51.81, the type strains of type species of the genera *Neisseria*, *Kingella*, *Eikenella*, and the type strains of five species in question for reclassification grouped them into three clusters (Fig. 3). On one hand, *Eikenella* cluster included *Neisseria shayeganii* strain 871^T^ along with type species *Eikenella corrodens* NCTC10586^T^ and other type strains of *Eikenella* species. On the other hand, *Neisseria* cluster consisted of *Uruburuella suis* DSM 17474^T^*, U. testudinis* CCUG 63373^T^, *Kingella potus NCTC 13336^T^, Bergeriella denitrificans* NCTC 10295^T^, *Morococcus cerebrosus* CIP 81.93^T^, strains 3986^T^ and 51.81 along with type species *Neisseria gonorrheae* DSM 9188^T^ and other type strains of *Neisseria* species. A threshold of 70.65% accurately separated these clusters except for *Neisseria shayeganii* strain 871^T^ which had an AAI higher than 70.65% with some *Neisseria* species. Nevertheless, the AAI values between *Neisseria shayeganii* strain 871^T^ and type species *Eikenella corrodens* NCTC10586^T^ were higher (74.41%) in comparison to that with *Neisseria gonorrheae* DSM 9188^T^ (69.67%). Considering the core-gene phylogeny, the DUS dialects found and the AAI calculations, we propose the reclassification of *Morococcus cerebrosus, Bergeriella denitrificans, Kingella potus, Uruburuella suis* and *Uruburuella testudinis* into *Neisseria* genus and *Neisseria shayeganii* into *Eikenella* genus to limit sources of confusion within the family Neisseriaceae.

**Fig. 3:**
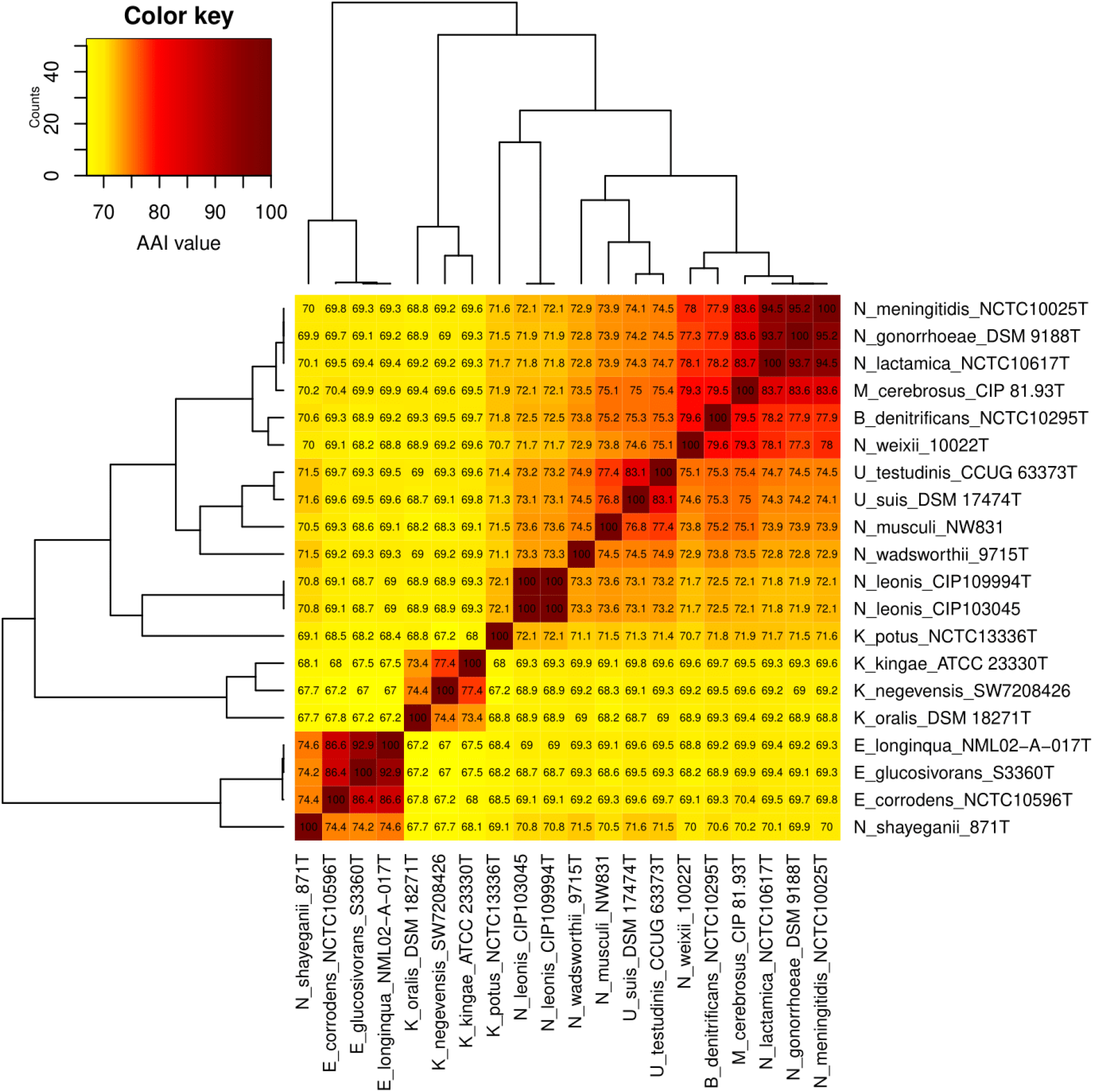
Pairwise AAI values between strains 3986^T^, 51.81 and representative strains of *Neisseria, Kingella, Morococcus, Bergeriella, Uruburuella* and *Eikenella*. The clustering between rows and columns was done with 1-Pearson correlation coefficient and UPGMA agglomeration method. Three distinct clusters appear from bottom-left to top-right for the genera *Eikenella*, *Kingella* and *Neisseria*.

The DUS dialect used in strains 3986^T^ and 51.81 was AG-DUS: it was repeated 1890 times and 1870 times while all other elements were repeated less than 1000 times (Table S4). The repetitive presence of DUS indicated that strains 3986^T^ and 51.81 were naturally competent to bind to and to take up exogenous DNA, as are all *Neisseria* strains. Like in most Gram-negative bacteria, *Neisseria* competence is possible via a Type IV Pili (Tfp) [37]. Tfp-like fibers are visible on the TEM image (Figs 4 and S5). For all *Neisseria* species, the *comP* pseudopilin on the bacterial surface can recognise environmental DNA fragments with a DUS motif [38]. The recognized DNA element binds to *comP* and enters the bacterium via the retraction of *comP* through the action of the *pilT* motor protein. From there, other machinery takes over to integrate the fragment into the bacterial DNA by transformation. Annotation of strains 3986^T^ and 51.81 genomes with the NCBI PGAP annotation module identified two coding sequences (CDS) related to the *pilT* protein and two others associated with a pseudopilin of the GspH/FimT family, whose role in bacterial transformation was highlighted this year [39]. The gene *comP* was not identified by PGAP: to verify its presence, a blastp was performed between the sequence A0A125WA94 of Uniprot associated with *Neisseria meningiditis* and the CDS identified by PGAP. There was a single hit for both strains: “pgaptmp_000082 pilin [*Neisseria*]” for strain 3986^T^ and “pgaptmp_001752 pilin [*Neisseria*]” for strain 51.81 achieved 96% coverage for 36.73% identity.

**Fig. 4:**
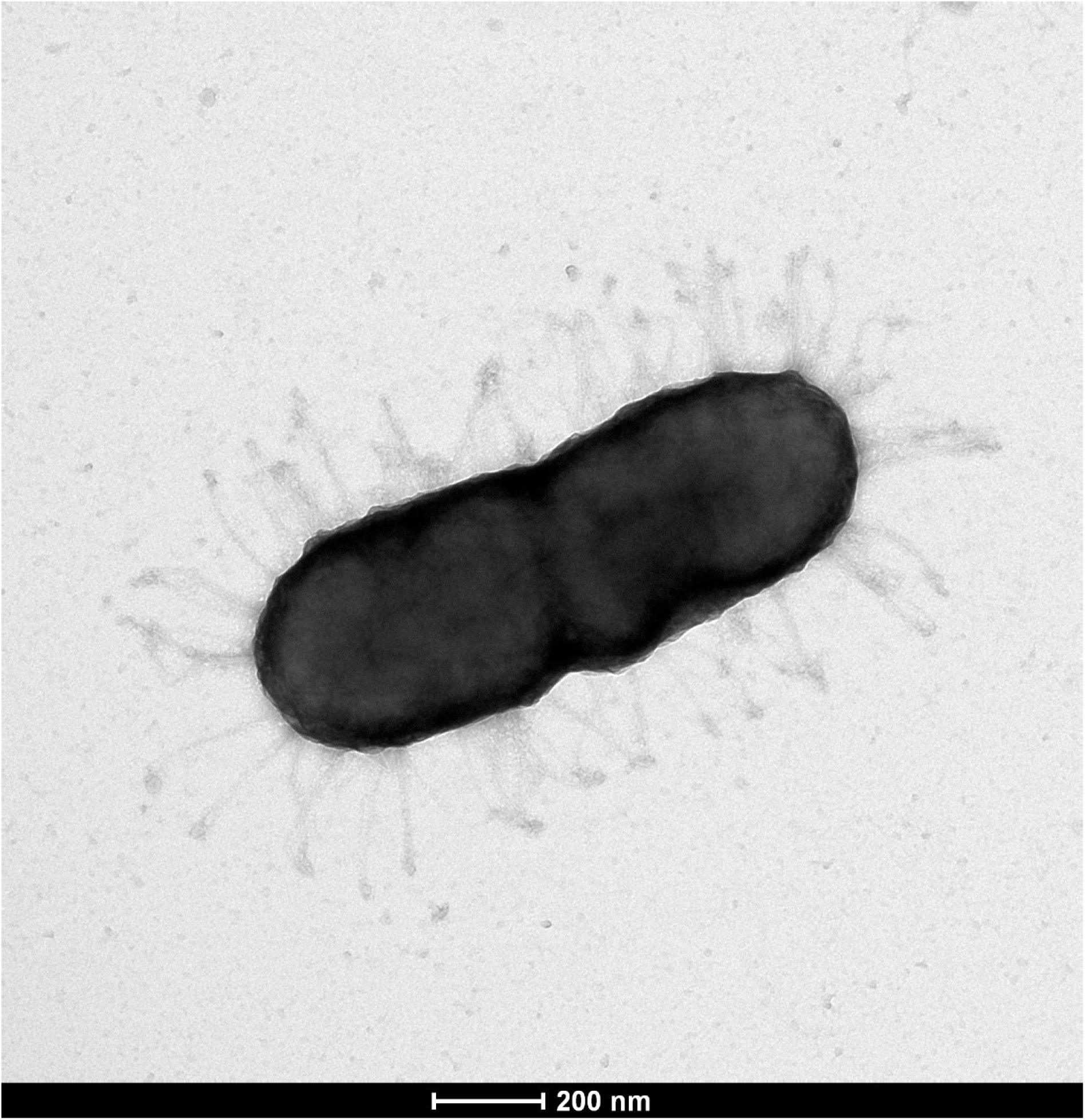
Transmission electron microscopy image of strain 3986^T^. The cell is a coccobacilli surrounded by Tfp-like fibers. Scale bar 200 nm.

All the strains of proposed new species including the strain 3986^T^ were isolated from rabbits. We did not have any information about the health of the rabbits, but it appeared that they were ill given that they were autopsied. Additionally, commensal species of *Neisseria* serve as a reservoir of resistance genes and virulence factors for the pathogenic *Neisseria* species [40,41]. This motivated the search of virulence factors and resistance genes present in the genomes of these strains. A total of 34 virulence factors were found in strain 3986^T^ and 35 in strain 51.81 of which 33 were common. Strain 3986^T^ also contained a Salmochelin from *Klebsiella* while strain 51.81 contained a Ferric enterobactin transport protein A/ferric-repressed protein B and a nitrate reductase from *Mycobacterium* (Table S5). This coincided with the phenotypic characteristics performed which proved only strain 51.81 could reduce nitrate (and nitrite). Nitrate and nitrite reduction by *Neisseria* is a very common feature, as it potentially allows bacterium to grow in anaerobic conditions using oxidized nitrogen compounds as alternative electron acceptors for energy production [42]. A large majority of virulence factors, 22 of strain 3986^T^ and 23 of strain 51.81, were found in *Neisseria* VFDB, while the others were reported from diverse bacterial genera. These virulence factors were essentially related to adhesion (10 including 4 associated with the type IV pili system to which *pilT* belongs), iron uptake (6 plus 1 for strain 51.81), stress adaptation (5). The details of the virulence factors found are presented in the supplementary material (Table S5). Contrary to virulence factors, no perfect or strict hits were found with antimicrobial resistance genes by CARD. Furthermore, four putative secondary metabolite clusters were identified in the genomes of both strains using anti-SMASH pipeline: hserlactone serves for communication between fungi and bacteria [43], resorcinol is used in a wide variety of products from rubber to antiseptic, arylpyolene is a pigment protecting the bacterium from reactive oxygen species [44], and terpene is a large class of natural products with a variety of roles in mediating antagonistic and beneficial interactions among organisms [45].

### Physiology

Strains 3986^T^ and 51.81 grew well on TSA, Columbia agar with 10% horse blood and BHI. Colonies of strains 3986^T^ and 51.81 on TSA appeared small, round, shiny and translucent (Fig. S6). Both strains formed aggregates in BHI broth, but strain 3986^T^ floated on the top of the broth forming a pellicle while strain 51.81 sinked to the bottom forming sediment. The cells appeared coccobacilli or diplococcobacilli under the microscope (Figs 4, S5 and S7), which indicates that the stains are between their evolutionary neighbors *Neisseria elongata* and *Neisseria bacilliformis* which are rods [46,47] and most *Neisseria* species which are cocci [48]. Cell size varied from 1 to 2 μm. The optimal temperature for growth on TSA was 37°C (range 28 to 45 °C) and the optimal pH was 8.5 (range 5.5 to 10 with only a slight growth at 5.5). Growth was equally fast with or without 5% CO^2^ and both strains could still grow with 0.5% oxgall added to TSB, which coincided with the discovery of one of the strains in the liver of a rabbit, where bile salts are highly concentrated. Growth occurred in microaerophilic conditions but not in anaerobic conditions. Strains 3986^T^ and 51.81 were gram-negative, non-motile and non-hemolytic. They were catalase and oxidase positive as most *Neisseria*, except for some strains of *Neisseria elongata* and *Neisseria bacilliformis* which are catalase negative [48].

The strains 3986^T^ and 51.81 showed indole production while the other strains DSM 23338^T^ (*N. bacilliformis*), CIP 108935^T^ (*K. potus*), CIP 106968^T^ (*N. dentiae*), CIP 72.27^T^ (*N. elongata*), and CIP 79.18^T^ (*N. gonorrheae*) did not produce indole (Table 1). Indole serves as intracellular, interspecies and even interkingdom signaling molecule [49]. Indole production appears a distinctive feature for strains 3986^T^ and 51.81, and blastn research of UniProt protein P0A853 (Tryptophanase of *Escherichia coli* strain K12) against the genomes dataset including all type strains of the genus *Neisseria* resulted only with the genomes of strains 3986^T^ and 51.81. API ZYM showed lipase C14 could be another distinctive feature (Table S6). Strain 51.81 was found to be nitrate and nitrite positive while strain 3986^T^ was not (Table 1). Antibiograms (Table S7) proved both 3986^T^ and 51.81 were contact resistant to lincomycin and vancomycin which is in agreement with intrinsic resistances of *Neisseria* species according to CA-SFM/EUCAST [50]. On the contrary, the intrisic resistance to thimethoprim was not found in any strain, while the strain 51.81 was contact resistant to streptomycin.

**Table 1:** Result of biochemical testing for *Neisseria leonis* sp. nov., in comparison with the phylogenetically closely related species from the genera *Neisseria* and *Kingella*. Strains: 1, 3986^T^; 2, 51.81; 3, DSM 23338^T^ (*N. bacilliformis);* 4, CIP 108935^T^ (*K. potus);* 5, CIP 106968^T^ (*N. dentiae);* 6, CIP 72.27^T^ (*N. elongata);* 7, CIP 79.18^T^ (*N. gonorrheae*).

| Characteristic | 1 | 2 | 3 | 4 | 5 | 6 | 7 |
| --- | --- | --- | --- | --- | --- | --- | --- |
| Glucose | - | - | + | - | + | - | + |
| Fructose | - | - | - | - | + | - | - |
| Maltose | - | - | - | - | - | - | - |
| Saccharose | - | - | - | - | + | - | - |
| Ornithine decarboxylase | - | - | - | - | - | - | - |
| Urease | - | - | - | - | - | - | - |
| Lipase | + | - | - | - | - | - | - |
| Alkaline phosphatase | - | - | - | - | - | - | - |
| Beta galactosidase | - | - | - | - | - | - | - |
| Proline arylamidase | - | + | + | - | - | + | + |
| Gamma glutamyl transferase | - | - | - | - | - | - | - |
| Indole | + | + | - | - | - | - | - |
| Nitrate reduction | - | + | + | + | + | - | ND |
| Nitrite reduction | - | + | + | + | + | + | ND |

All data from this study. ND: No data

Based on the genetic and phenotypic characteristics, we propose that strains 3986^T^, 51.81 and CCUG 45853 are classified as members of a novel species, for which we propose the name *Neisseria leonis* sp. nov. with 3986^T^ as type strain. Additionally, based on core-gene phylogeny, dominant DUS dialect and AAI calculations, we propose the reclassification of *Morococcus cerebrosus, Bergeriella denitrificans, Kingella potus, Uruburuella suis* and *Uruburuella testudinis* to *Neisseria* genus and conversely the reattribution of *Neisseria shayeganii* to *Eikenella* genus.

## Description of *Neisseria leonis* sp. nov

*Neisseria leonis* (le.on’is N.L. gen. n. leonis of Léon, in honour of Léon Boutroux (1851-1921), a French chemist who worked under the direction of Louis Pasteur on glucose fermentation. He notably published a book on pan fermentation in 1897).

Cultures grow well on Trypcase Soy Agar, Columbia agar with 10% horse blood or Brain Heart Infusion. Optimal growth temperature is at 37°C and pH 8.5 and can tolerate bile salts at least up to 0.5%. Growth with or without 5% CO^2^ and in microaerophilic conditions. No growth in anaerobic conditions. Colonies are small, round, shiny and translucent. Cells are diplococcobacilli with length around 1-2 μm. Gram negative, non-motile, non-hemolytic, oxidase and catalase positive. Positive for production of indole, alkaline phosphatase, esterase (C4), esterase lipase (C8), lipase (C14), leucine arylamidase, valine arylamidase, acid phosphatase and naphtol-AS-BI-phosphohydrolase. Negative for production of ornithine decarboxylase, urease, alkaline phosphatase, ß-galactosidase, gamma-glutamyltransferase, trypsin, alpha-chymotrypsin, alpha-galactosidase, ß-glucuronidase, alpha-glucosidase, ß-glucosidase, N-acétyl-ß-glucosaminidase, alpha-mannosidase and alpha-fucosidase. Negative for utilization of D-glucose, D-fructose, D-maltose and D-saccharose. Contact resistance with the antibiotics lincomycin and vancomycin. The DNA G+C content of the type strain is 56.92%.

Type strain of the species is 3986^T^ (= R726^T^ = CIP 109994^T^ = LMG 32907^T^). Type strain was isolated from the liver of a rabbit in 1972 in the Institut Pasteur of Lyon, France. The GenBank sequence accession number of the genome sequence is JAPQFK000000000 and the 16S rRNA gene sequence of is OQ121838.1.

## Description of *Neisseria cerebrosus* comb. nov

*Neisseria cerebrosus* (ce.re.bro.sus. L. masc. adj. *cerebrosus*, having a madness of the brain, hare-brained, hotbrained, passionate, intended to mean pertaining to the brain, the original source of isolation of this organism)

Basonym: *Morococcus cerebrosus* [56]

The description is the same as for *M. cerebrosus* [56]. Phylogenetic analysis with the *bac120* gene set from GTDB, repetitive presence of one of the *Neisseria* DUS dialect and AAI values provided strong evidence for the placement of this species in the genus *Neisseria*. The type strain is UQM 858^T^ (= ATCC 33486^T^ = NCTC 11393^T^).

## Description of *Neisseria denitrificans* comb. nov

*Neisseria denitrificans* (de.ni.tri’fi.cans. L.prep. de away from; L. nitrum soda; N. L. n. nitrum nitrate; N. L. v. denitrifico to denitrify; N. L. part. adj. denitrificans denitrifying)

Basonym: *Bergeriella denitrificans* [54]

The description is the same as for *N. denitrificans* [55]. It was proposed to reclassify this strain as the type species of a new genus *Bergeriella* based on 16S phylogeny [54] but modern genomic analysis based on phylogenetic analysis with the *bac120* gene set from GTDB, repetitive presence of one of the *Neisseria* DUS dialect and AAI values provided strong evidence for the placement of this species in the genus *Neisseria*. The type strain is ATCC 14686^T^ (= CCUG 2155^T^ = NCTC 10295^T^)

## Description of *Neisseria potus* comb. nov

*Neisseriapotus* (po.tus. L. gen. masc. n. *potus*, of the drink or drinking, pertaining to *Potus flavus*, the generic name of the South American kinkajou, the animal from which the organism originated)

Basonym: *Kingella potus* [53]

The description is the same as for *K. potus* [53]. Phylogenetic analysis with the *bac120* gene set from GTDB, repetitive presence of one of the *Neisseria* DUS dialect and AAI values provided strong evidence for the placement of this species in the genus *Neisseria*. The type strain is 3/SID/1128^T^ (= NCTC 13336^T^ = CCUG 49773^T^).

## Description of *Neisseria suis* comb. nov

*Neisseria suis* (su’is. L. fem. n. sus, suis pig, hog; L. gen. n. suis of the hog)

Basonym: *Uruburuella suis* [51]

The description is the same as for *U. suis* [51]. Phylogenetic analysis with the *bac120* gene set from GTDB, repetitive presence of one of the *Neisseria* DUS dialect and AAI values provided strong evidence for the placement of this species in the genus *Neisseria*. The type strain is 258/02^T^ (= CCUG 47806^T^ = CECT 5685^T^).

## Description of *Neisseria testudinis* comb. nov

*Neisseria testudinis* (tes.tu’di.nis. L. n. testudo tortoise; L. gen. n. testudinis of the tortoise)

Basonym: *Uruburuella testudinis* [52]

The description is the same as for *U. testudinis* [52]. Phylogenetic analysis with the *bac120* gene set from GTDB, repetitive presence of one of the *Neisseria* DUS dialect and AAI values provided strong evidence for the placement of this species in the genus *Neisseria*. The type strain is 07_OD624^T^ (= DSM 26510^T^ = CCUG 63373^T^).

## Description of *Eikenella shayeganii* comb. nov

*Eikenalla shayeganii* (sha.ye.ga’ni.i. N.L. gen. n. shayeganii of Shayegani, to recognize and honour over 40 years of public service and leadership by Dr Mehdi Shayegani in the Bacteriology Laboratory of the Wadsworth Center, New York State Department of Health)

Basonym: *Neisseria shayeganii* [57]

The description is the same as for *N. shayeganii* [57]. Phylogenetic analysis with the *bac120* gene set from GTDB, repetitive presence of the *Eikenella* DUS dialect AG-eikDUS and AAI values provided strong evidence for the placement of this species in the genus *Eikenella*. The type strain is 871^T^ (= DSM 22246^T^ = CIP 109933^T^).

## Abbreviations

AAI: average amino acid identity
ANI: average nucleotide identity
ASV: Amplicon Sequence Variant
CDS: coding sequences
CIP: Collection of Institut Pasteur
dDDH: digital DNA-DNA hybridization values
DUS: DNA uptake sequence
GTDB: Genome Taxonomy Database
ICNP: International Code of Nomenclature of Prokaryotes
LPSN: List of Prokaryotic names with Standing in Nomenclature
ML: maximum likelihood
NJ: neighbor joining
MP: maximum parsimony
TEM: Transmission Electron Microscopy
TSA: trypticase soy agar
TSB: tryptone soy broth
VFDB: virulence factor database.

## Acknowledgements

We are grateful to our colleagues of the DSMZ (German Collection of Microorganisms and Cell Cultures) for their gift of strain DSM 23338^T^.

