## Supplementary texts, figures and legends of supplementary tables for "*Neisseria leonis* sp. nov. isolated from rabbits, reclassification of *Uruburuella suis, Uruburuella testudinis, Kingella potus, Bergeriella denitrificans* and *Morococcus cerebrosus* into *Neisseria* genus and reclassification of *Neisseria shayeganii* into *Eikenella* genus"

**Title:**

**Authors and affiliations:**

M. Boutroux[1], S. Favre-Rochex[1], O. Gorgette[2], G. Touak[1], E. Muhle[1], O. Chesneau[1], D. Clermont[1], P. Rahi[1]

[1] Institut Pasteur, Université Paris Cité, Collection of Institut Pasteur (CIP), F-75015 Paris, France
[2] Institut Pasteur, Université Paris Cité, Ultrastructural BioImaging Platform (UBI), F-75015 Paris, France

**Corresponding authors details:**

Martin Boutroux,
Collection de l'Institut Pasteur, Paris, France


Praveen Rahi,
Collection de l'Institut Pasteur, Paris, France


**Supplementary text 1:** Metagenomic analysis between strain $3986^T$ with public microbiome rabbit datasets

*Material and methods:*

Rabbit microbiome data from a 2019 study [1] was searched to find Amplicon Sequence Variants (ASVs) identical to strain $3986^T$. Data was loaded on QIIME2 2022.11 [2] with the q2-fondue 2022.2 plugin [3]. The appropriate trimming parameters were calculated with FIGARO 1.1.2 [4] and used in DADA2 [5] to produce the Amplicon Sequence Variant (ASV) table. A BLAST was calculated between these ASVs and the V4 region extracted from the 16S rRNA sequence of strain $3986^T$.

*Results:*

An ASV present 19 times in one of the 126 samples (SRR6475038) is 100% identical to the sequence of strain $3986^T$. The characteristics of this ASV were:

**Run :** SRR6475038 from SRX3564915

**BioSample :** SAMN08358139

| | |
|---|---|
| **host** | Oryctolagus cuniculus |
| **collection date** | 2015-11-06 |
| **isolation source** | feces |
| **geographic location** | Spain:Rabanales, Cordoba |
| **latitude and longitude** | missing |
| **sex** | female |
| **age** | > 2 years |
| **host_subsp** | cuniculus |
| **individual** | P.44 |
| **origin** | P |
| **plot** | 6 |
| **warren** | 8 |
| **cage** | NA |
| **weight** | 1374 |
| **tarsus** | 61.74 |
| **fate** | alive |

**Sequence:**

TACGTAGGGTGCGAGCGTTAATCGGAATTACTGGGCGTAAAGCGGGCGCAGACGGTTTGTTAAG
CAGGATGTGAAATCCCCGGGCTCAACCTGGGAACTGCGTTCTGAACTGGCAGGCTAGAGTGTGT
CAGAGGGGGGTAGAATTCCACGTGTAGCAGTGAAATGCGTAGAGATGTGGAGGAATACCGATGG
CGAAGGCAGCCCCCTGGGATAACACTGACGTTCATGCCCGAAAGCGTGGGTAGCAAACAGG

A similar search for identical sequences in the fecal microbiome of different herbivores among which rabbits (SRP405849) (no publication associated yet) resulted in zero BLAST hit, confirming the rarity of the new *Neisseria* species.

*References :*

**Supplementary text 2:** Characteristics of the dataset used for the phylogenetic analysis of the whole genome sequences

*Methods:*

The dataset comprised 79 genomes including genomes of strain 3986$^T$, strain 51.81 and *Chromobacterium violaceum* used as an outgroup. There was the 31 *Neisseria* species validated on LPSN (3 of these sequences are not those of the type strain) completed by 2 species not validated on LPSN. Bacterial genera close to *Neisseria* were identified with the MEGA 11 Timetree tool and added to the dataset, namely the unique species of the genera *Bergeriella* and *Simonsiella,* the 6 species of *Kingella* (5 validated on LPSN including one genome from a non-type strain and 1 non-validated species), the 2 species of *Uruburuella*. The genomes of the species returned in the EZBioCloud results as well as the type species of the associated genera were also integrated into the dataset. The additional genomes were 23 *Neisseria* species according to GTDB but which have not been described. The information concerning the dataset is available in the Excel file Table_S3_caracs_genomes_references.xlsx. It contains metadata (accession number, organism name, isolate, type strain or not), quality metrics determined with contig_info (number of contigs, genome size, %GC, N50) as well as a verification of the purity of the genomes with checkM (completeness, contamination, strain heterogeneity). LPSN included 1 additional *Neisseria* species (*Neisseria oralis*) as well as 2 non-validated *Neisseria* species (*"Neisseria kochii"* and *"Neisseria skkuensis"*) but no genomes were available for these species. A blastn of the 16S of our two strains with the 16S of these 3 species (corresponding respectively to accession numbers NR_118249.1, U02900.1, FJ763637.1) did not return any significant alignment. Moreover, species *Kingella pumchi* was described too recently to be included in the analysis [1] but an ANI calculation between GCA_022288825.1 and the genome of strain 3986$^T$ showed a limited 78.66 % identity.

*References:*

1. Xiao M, Liu R, Du J, Liu R, Zhai L, Wang H, et al. Kingella pumchi sp. nov., an organism isolated from human vertebral puncture tissue. Antonie Van Leeuwenhoek. 2023 Feb;116(2):143–51

**Table S1:** List of EZBioCloud hits for strain 3986$^{T}$ 16S rRNA sequence

(see Excel file Table_S1_EZBioCloud_16S_hits_for_3986T.xlsx)

**Table S2:** List of TYGS hits with the genome of strain 3986$^{T}$

(see Excel file Table_S2_TYGS_job_results.xlsx)

**Table S3:** Characteristics of the genomes used for the phylogenetic analysis

(see Excel file Table_S3_caracs_genomes_references.xlsx)

**Table S4:** Count of each of the 8 DNA Uptake Sequences among the 79 whole genomes of the dataset with Jellyfish

(see Excel file Table_S4_DUS_counts.xlsx)

**Table S5:** Virulence factors of strains 3986$^{T}$ and 51.81 resulted by VFAnalyzer

(see Excel file Table_S5_virulence_factors_found.xlsx)

**Table S6:** API ZYM results for strains 3986$^{T}$ and 51.81 as well as the reference strains (including *Neisseria gonorrhoeae*)

(see Excel file Table_S6_API_ZYM_results.xlsx)

**Table S7:** Results of disk diffusion assay for strains 3986$^{T}$ and 51.81 as well as the reference strains

(see Excel file Table_S7_antibiograms_results.xlsx)

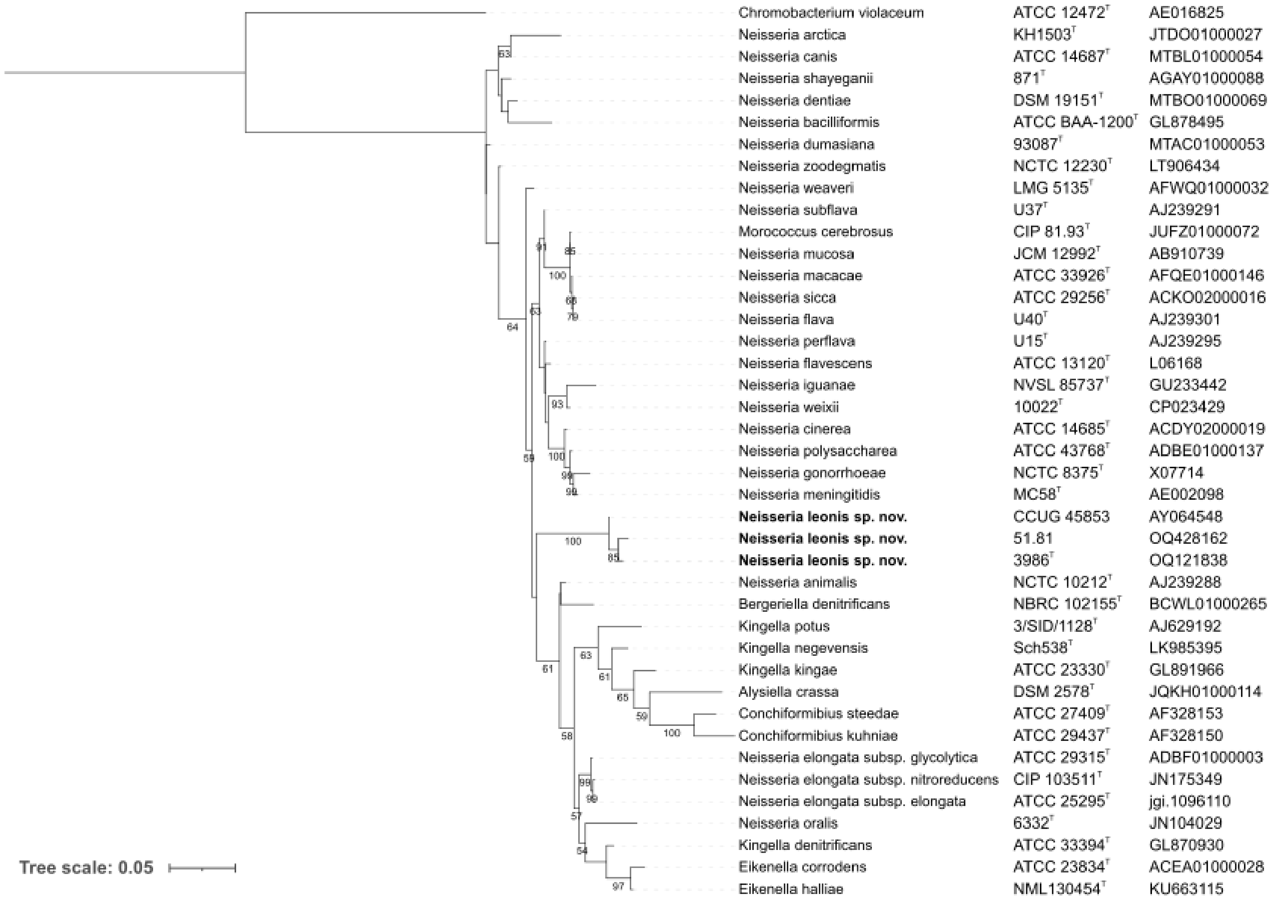


**Fig. S1:** 16S phylogenetic tree obtained with maximum likelihood method from the sequences of strains $3986^T$, 51.81, CCUG 45853 and 37 hits obtained in EZBioCloud with strain $3986^T$. TPM3+F+I+G4 model chosen according to BIC by ModelFinder. 1000 ultrafast bootstrap replications were performed. *Chromobacterium violaceum* ATCC $12472^T$ was used as an outgroup. Bootstrap values above 50% are displayed. Bar, 0.01 changes per site.

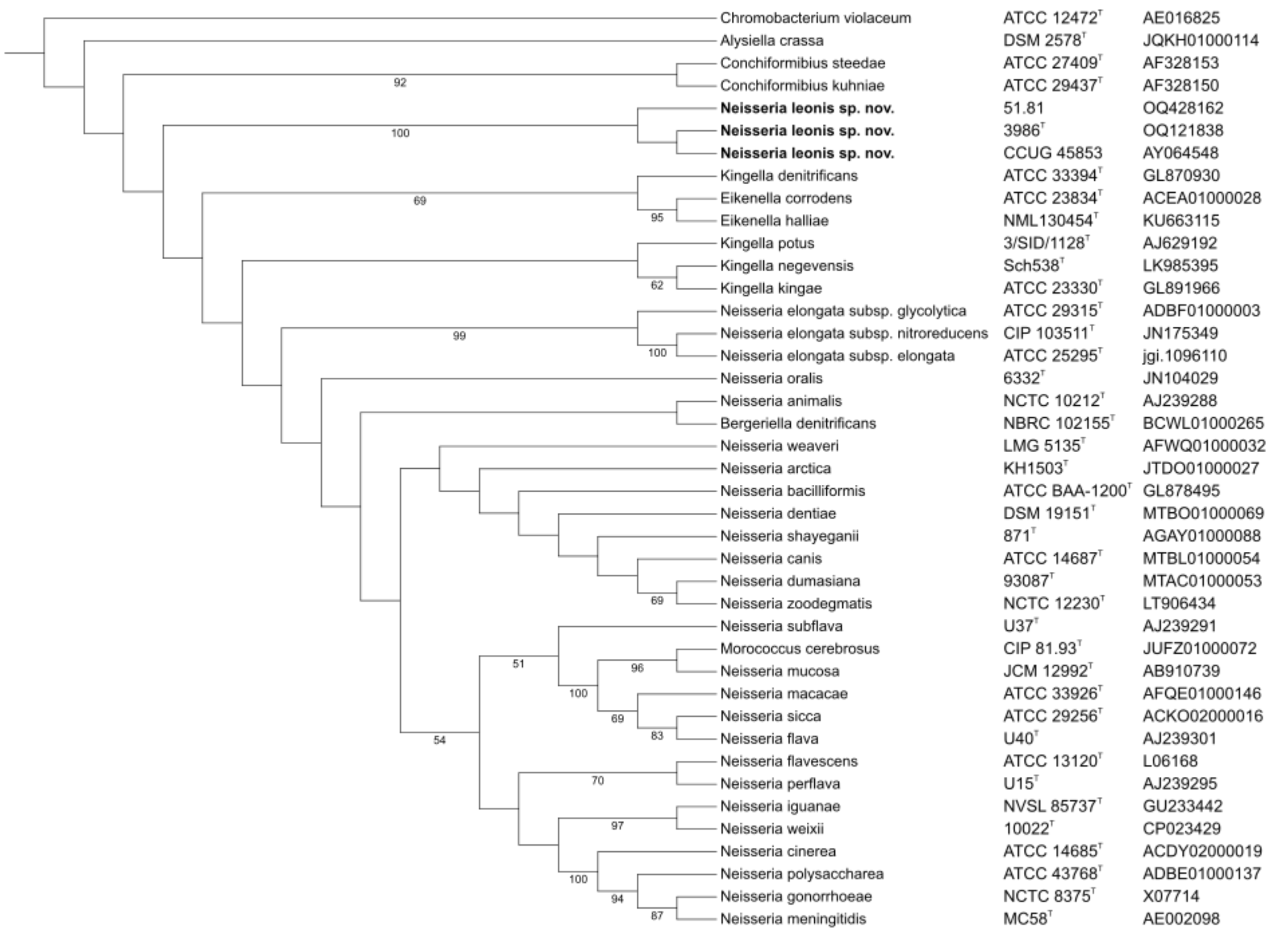


**Fig. S2:** 16S phylogenetic tree obtained in MEGA 11 with maximum parsimony method from the sequences of strains 3986$^{T}$, 51.81, CCUG 45853 and 37 hits obtained in EZBioCloud with strain 3986$^{T}$. *Chromobacterium violaceum* ATCC 12472$^{T}$ was used as an outgroup. Bootstrap values above 50% are displayed.

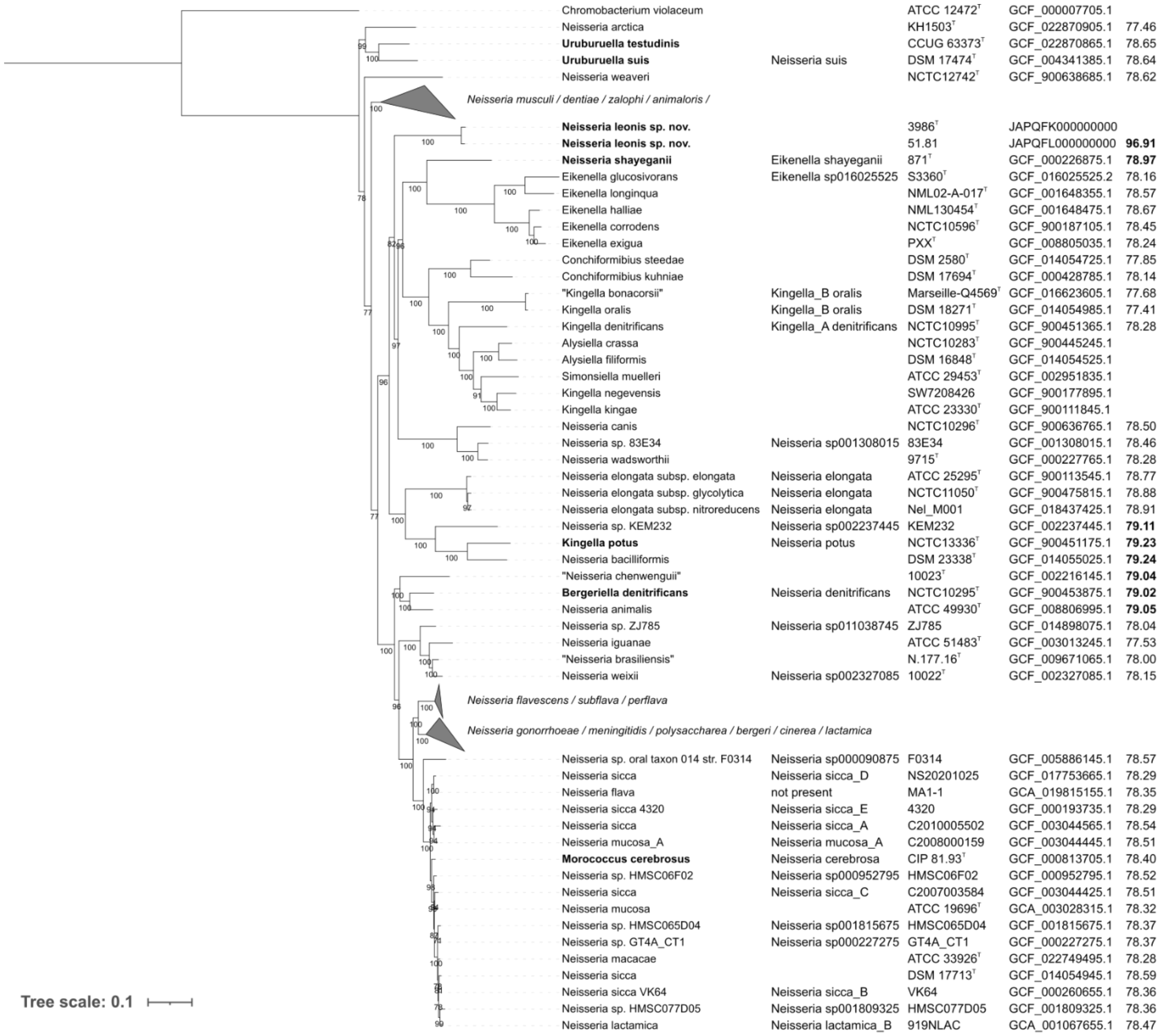


**Fig. S3:** Phylogenetic tree obtained from the alignment of the 53 rMLST genes with maximum likelihood method. Each line contains the name of the species according to LPSN and according to GTDB if it differs, the isolate, the accession number and the highest ANI values with strain 3986$^T$ (top 10 values are in bold). Three *Neisseria* clades are collapsed here because they are not relevant for the species that concerned us. GTR+F+R6 model chosen according to BIC by ModelFinder. 1000 ultrafast bootstrap replications were performed. *Chromobacterium violaceum* ATCC 12472$^T$ was used as an outgroup. Bootstrap values above 50% are displayed. Bar, 0.1 changes per site.

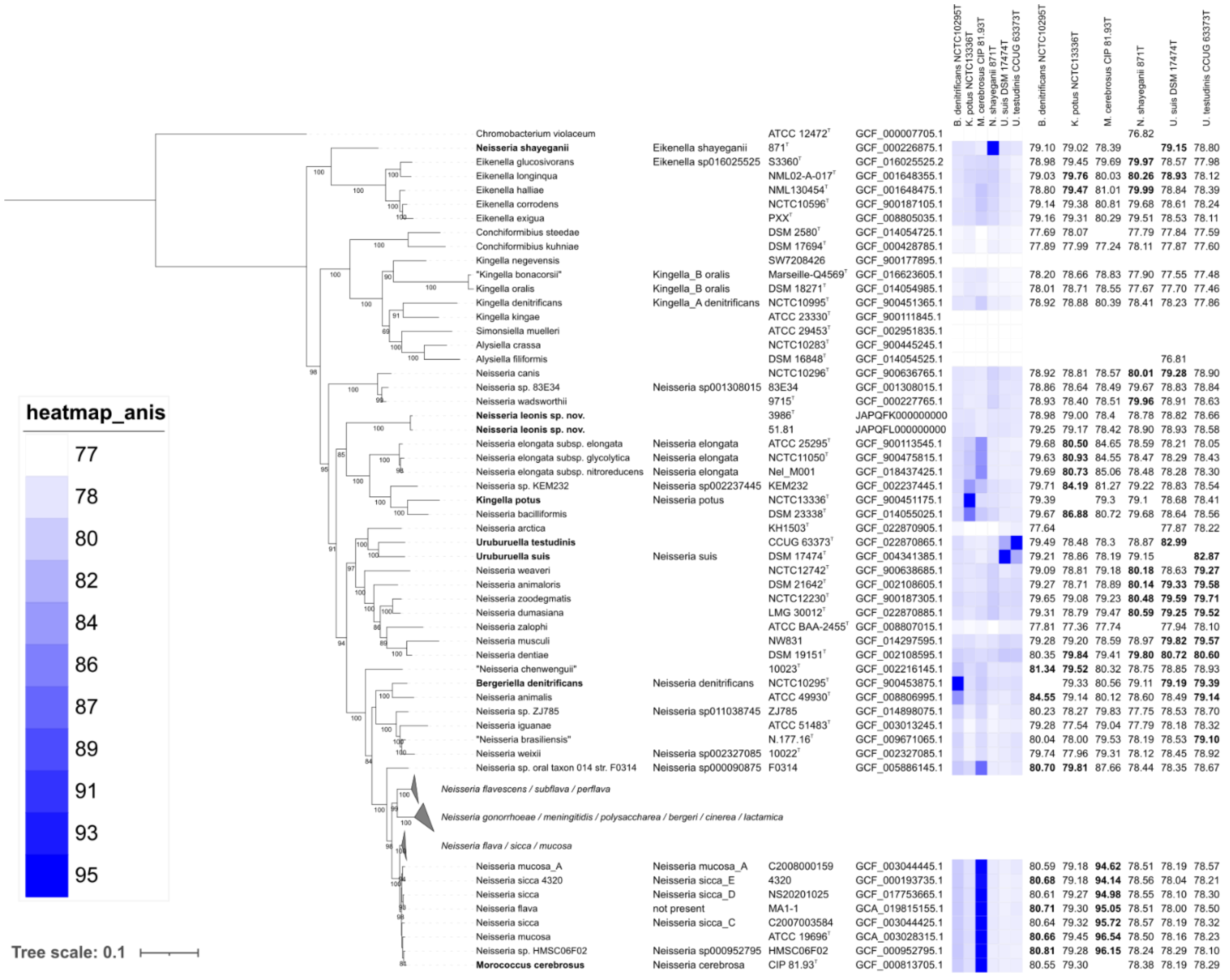


**Fig. S4:** Phylogenetic tree obtained from the alignment of the *bac120* genes with maximum likelihood method. Each line contains the name of the species according to LPSN and according to GTDB if it differs, the isolate name and the accession number. The heatmap depicts the highest ANI between the genomes of the dataset and the 6 misclassified species *Bergeriella denitrificans* NCTC10295$^{T}$, *Kingella potus* NCTC13336$^{T}$, *Morococcus cerebrosus* CIP 81.93$^{T}$, *Neisseria shayeganii* 871$^{T}$, *Uruburuella suis* DSM 17474$^{T}$ and *Uruburuella testudinis* CCUG 63373$^{T}$. The subsequent columns show the highest ANI values (top 10 values are in bold). Three *Neisseria* clades are collapsed here because they are not relevant for the species that concerned us. *Chromobacterium violaceum* ATCC 12472$^{T}$ was used as an outgroup. Bootstrap values above 50% are displayed. Bar, 0.1 changes per site.

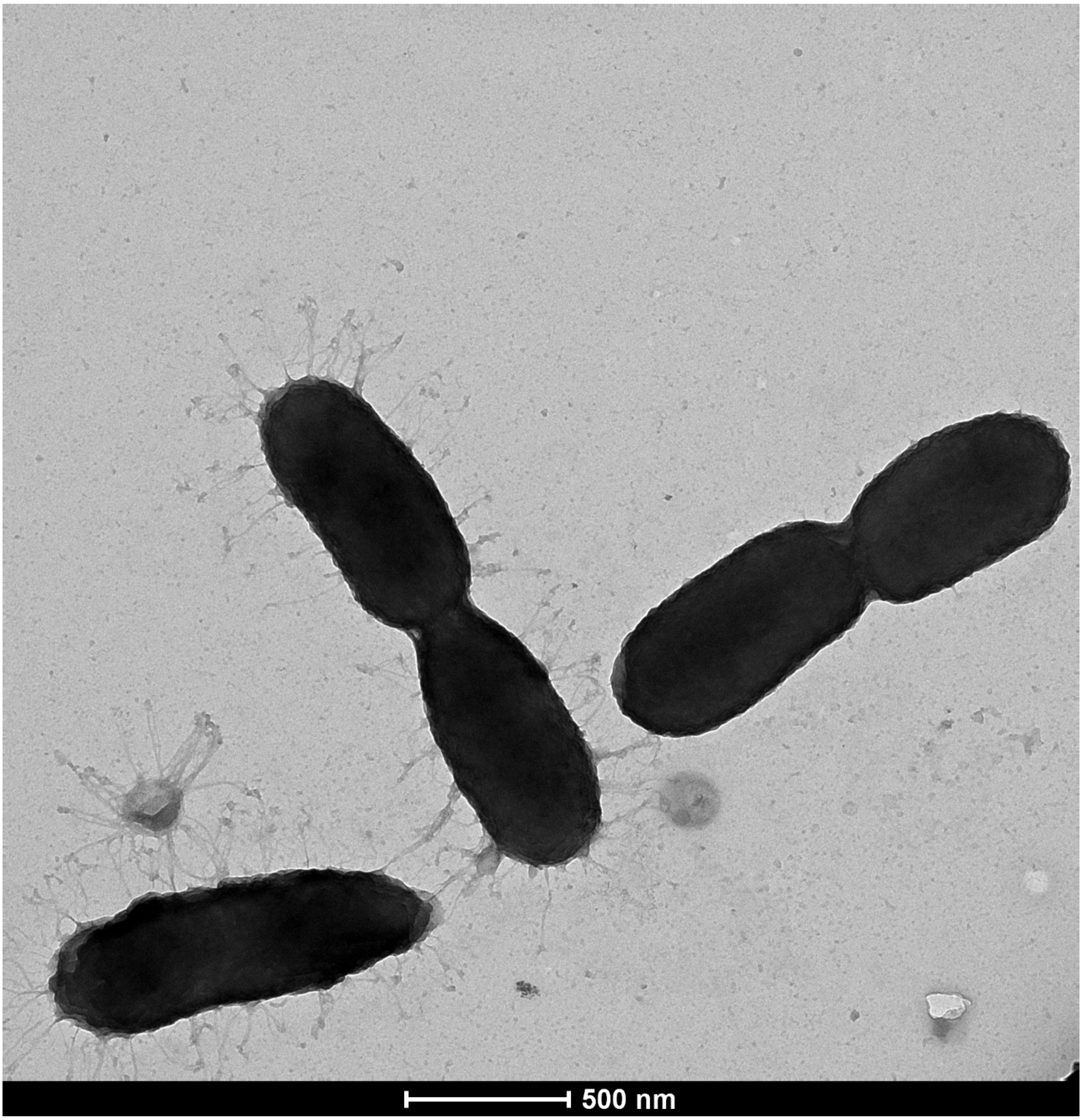


**Fig. S5:** Transmission electron microscopy image of strain 3986$^{T}$. Two pairs of diplococcobacilli and one coccobacilli appears connected by Tfp-like fibers. Scale bar 500 nm.

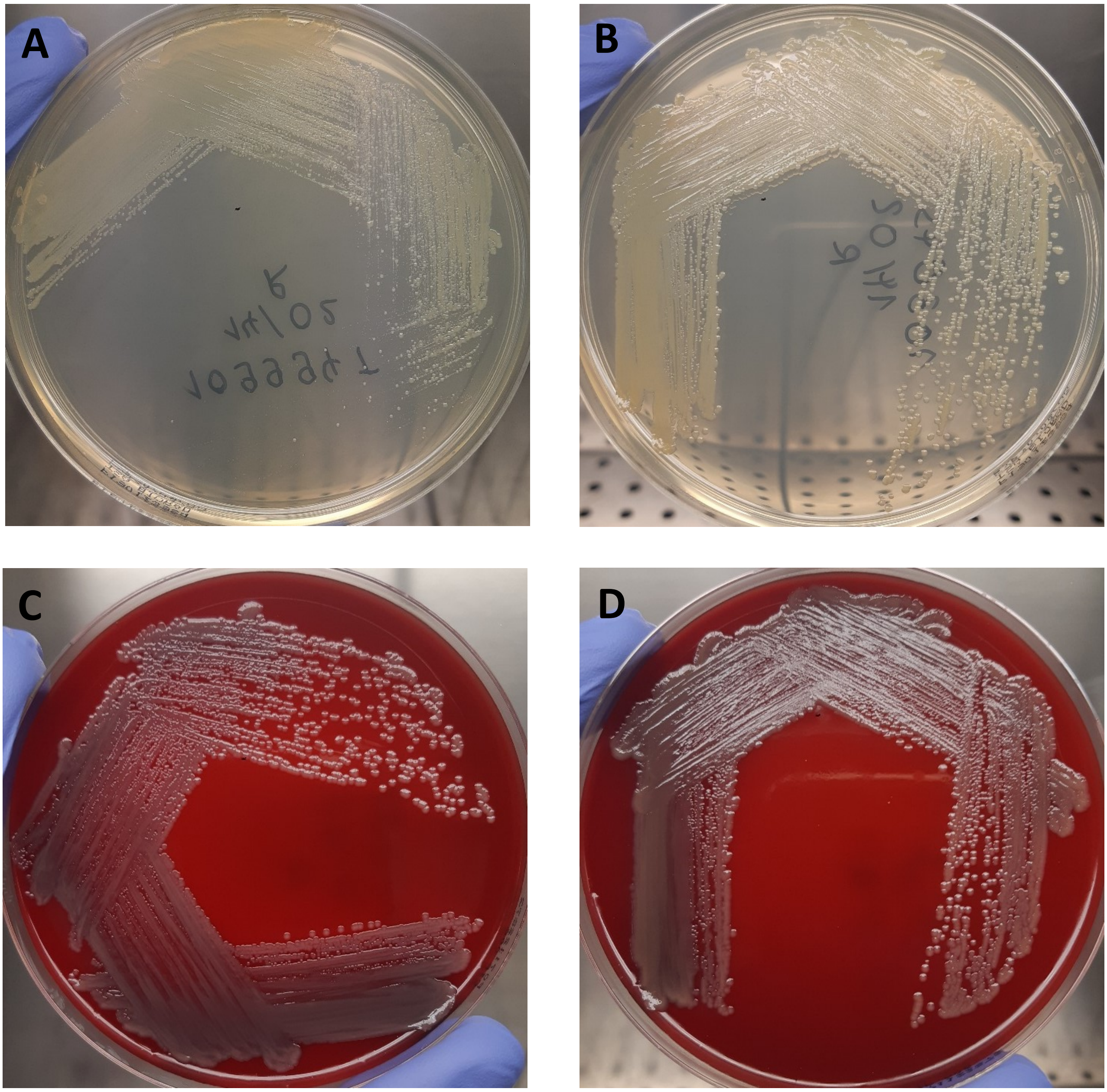


**Fig. S6:** Colonies of strains $3896^T$ and 51.81 after 24 hrs incubation at 37°C as observed on (a, b) Trypticase Soy Agar; (c, d) Columbia agar with 10 % horse blood

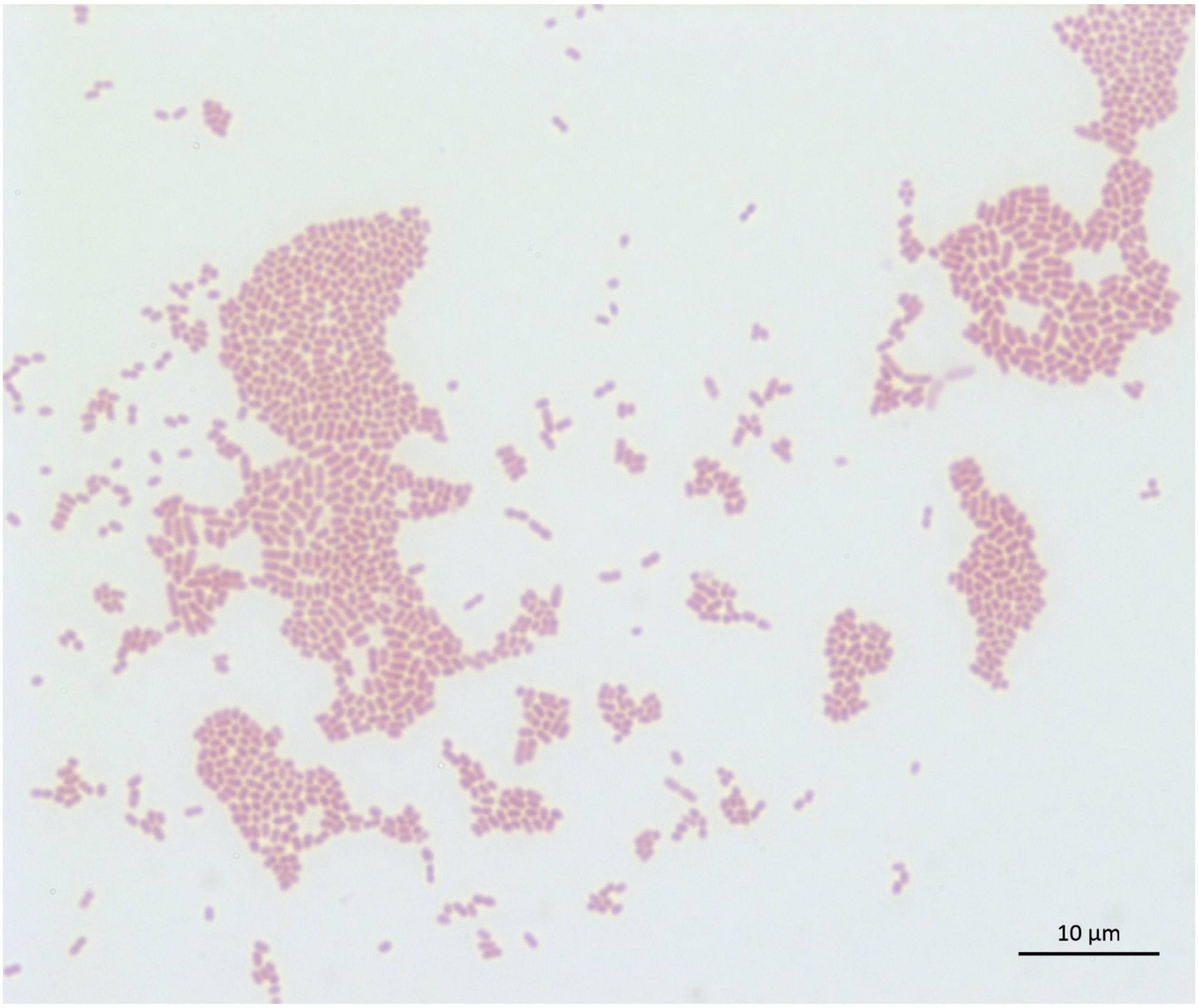


**Fig. S7:** Image obtained under an optical microscope at magnification 100X of the strain 3986$^T$. Cells are Gram-negative and form clear aggregates. Scale bar 10 µm.
