## Supplementary table 2 for "*Neisseria leonis* sp. nov. isolated from rabbits, reclassification of *Uruburuella suis, Uruburuella testudinis, Kingella potus, Bergeriella denitrificans* and *Morococcus cerebrosus* into *Neisseria* genus and reclassification of *Neisseria shayeganii* into *Eikenella* genus"

### TYPE (STRAIN) GENOME SERVER

### JOB SUMMARY

PRINT DATE: 2022-09-22 13:45:18 +0200

JOB ID: 0ebc395f-aa55-451b-bcc9-72b8ea216fdb

RESULT PAGE: https://tygs.dsmz.de/user_results/show?guid=0ebc395f-aa55-451b-bcc9-72b8ea216fdb

#### Table 1: Phylogenies

**Publication-ready versions** of both the genome-scale GBDP tree and the 16S rRNA gene sequence tree can be customized and exported either in SVG (vector graphic) or PNG format from within the phylogeny viewers in your TYGS result page. For publications the **SVG format is recommended** because it is lossless, always keeps its high resolution and can also be easily converted to other popular formats such as PDF or EPS. Please follow the link provided above!

#### Table 2: Identification

The below list contains the result of the TYGS species identification routine.

Explanation of remarks that might occur in the below table:

**remark [R1]:** The TYGS type strain database is automatically updated on an almost daily basis. However, if a particular type strain genome is not available in the TYGS database, this can have several reasons which are detailed in the FAQ. You can request an extended 16S rRNA gene analysis via the 16S tree viewer found in your result page to detect **not yet genome-sequenced** type strains relevant for your study.

**remark [R2]:** > 70% dDDH value (formula $d_4$) and (almost) minimal dDDH values for gene-content formulae $d_0$ and $d_6$ indicate a potentially unreliable identification result and should thus be checked via the 16S rRNA gene sequence similarity. Such strong deviations can, in principle, be caused by sequence contamination.

**remark [R3]:** G+C content difference of > 1 % indicates a potentially unreliable identification result because within species G+C content varies no more than 1 %, if computed from genome sequences (PMID: 24505073).

| Strain | Conclusion | Identification result | Remark |
|---|---|---|---|
| **'CIP109994.scf'** | potential new species | | see [R1] |
| **'CIP103045.scf'** | potential new species | | see [R1] |

#### Table 3: Pairwise comparisons of user genomes vs. type-strain genomes

The following table contains the pairwise dDDH values between your user genomes and the selected type-strain genomes. The dDDH values are provided along with their confidence intervals (C.I.) for the three different GBDP formulas:

- formula $d_0$ (a.k.a. GGDC formula 1): length of all HSPs divided by total genome length
- formula $d_4$ (a.k.a. GGDC formula 2): sum of all identities found in HSPs divided by overall HSP length
- formula $d_6$ (a.k.a. GGDC formula 3): sum of all identities found in HSPs divided by total genome length

**Note:** Formula $d_4$ is independent of genome length and is thus robust against the use of incomplete draft genomes. For other reasons for preferring formula $d_4$, see the FAQ.

| Query | Subject | $d_0$ | C.I. $d_0$ | $d_4$ | C.I. $d_4$ | $d_6$ | C.I. $d_6$ | Diff. G+C Percent |
|---|---|---|---|---|---|---|---|---|
| **'CIP109994.scf.fasta'** | **'CIP103045.scf.fasta'** | 88.0 | [84.5 - 90.8] | 70.8 | [67.7 - 73.6] | 87.8 | [84.8 - 90.3] | 0.03 |
| **'CIP109994.scf.fasta'** | *Neisseria dentiae* DSM 19151 | 17.6 | [14.6 - 21.2] | 23.8 | [21.5 - 26.3] | 17.6 | [15.0 - 20.6] | 2.52 |
| **'CIP103045.scf.fasta'** | *Neisseria dentiae* DSM 19151 | 17.6 | [14.6 - 21.2] | 23.7 | [21.4 - 26.1] | 17.6 | [14.9 - 20.5] | 2.55 |
| **'CIP109994.scf.fasta'** | *Neisseria chenwenguii* 10023T | 15.2 | [12.3 - 18.6] | 22.8 | [20.5 - 25.2] | 15.3 | [12.8 - 18.2] | 2.9 |
| **'CIP103045.scf.fasta'** | *Neisseria chenwenguii* 10023T | 15.3 | [12.4 - 18.8] | 22.7 | [20.4 - 25.1] | 15.5 | [13.0 - 18.4] | 2.93 |
| **'CIP103045.scf.fasta'** | *Neisseria animalis* DSM 23392 | 15.4 | [12.5 - 18.9] | 22.7 | [20.4 - 25.1] | 15.6 | [13.0 - 18.5] | 5.09 |
| **'CIP109994.scf.fasta'** | *Neisseria mucosa* ATCC 19696 | 14.9 | [12.1 - 18.4] | 22.6 | [20.3 - 25.0] | 15.1 | [12.6 - 18.0] | 5.83 |
| **'CIP103045.scf.fasta'** | *Bergeriella denitrificans* NBRC 102155 | 16.1 | [13.1 - 19.6] | 22.6 | [20.3 - 25.0] | 16.2 | [13.6 - 19.1] | 1.13 |
| **'CIP103045.scf.fasta'** | *Neisseria weixii* DSM 103441 | 14.4 | [11.6 - 17.8] | 22.5 | [20.2 - 24.9] | 14.7 | [12.2 - 17.5] | 7.74 |
| **'CIP103045.scf.fasta'** | *Kingella potus* DSM 18304 | 15.7 | [12.8 - 19.2] | 22.5 | [20.2 - 24.9] | 15.8 | [13.3 - 18.7] | 0.92 |
| **'CIP109994.scf.fasta'** | *Neisseria weixii* DSM 103441 | 14.7 | [11.8 - 18.1] | 22.4 | [20.1 - 24.9] | 14.9 | [12.4 - 17.7] | 7.7 |
| **'CIP103045.scf.fasta'** | *Neisseria mucosa* ATCC 19696 | 14.8 | [12.0 - 18.3] | 22.4 | [20.1 - 24.9] | 15.0 | [12.6 - 17.9] | 5.86 |
| **'CIP109994.scf.fasta'** | *Kingella potus* DSM 18304 | 15.9 | [13.0 - 19.4] | 22.4 | [20.2 - 24.9] | 16.0 | [13.5 - 18.9] | 0.95 |
| **'CIP109994.scf.fasta'** | *Neisseria animalis* DSM 23392 | 15.5 | [12.6 - 19.0] | 22.4 | [20.2 - 24.9] | 15.6 | [13.1 - 18.5] | 5.06 |
| **'CIP103045.scf.fasta'** | *Eikenella exigua* PXX | 14.2 | [11.4 - 17.6] | 22.4 | [20.2 - 24.9] | 14.5 | [12.0 - 17.3] | 1.53 |
| **'CIP109994.scf.fasta'** | *Neisseria zoodegmatis* DSM 21643 | 15.3 | [12.4 - 18.8] | 22.4 | [20.1 - 24.8] | 15.5 | [13.0 - 18.4] | 6.0 |
| **'CIP109994.scf.fasta'** | *Bergeriella denitrificans* NBRC 102155 | 16.2 | [13.3 - 19.7] | 22.4 | [20.1 - 24.8] | 16.3 | [13.7 - 19.2] | 1.1 |
| **'CIP103045.scf.fasta'** | *Neisseria bacilliformis* ATCC BAA-1200 | 15.4 | [12.5 - 18.9] | 22.3 | [20.0 - 24.7] | 15.6 | [13.1 - 18.5] | 2.62 |
| **'CIP103045.scf.fasta'** | *Neisseria zoodegmatis* DSM 21643 | 15.4 | [12.5 - 18.9] | 22.3 | [20.0 - 24.8] | 15.5 | [13.0 - 18.4] | 6.03 |
| **'CIP109994.scf.fasta'** | *Neisseria macacae* ATCC 33926 | 14.7 | [11.9 - 18.1] | 22.2 | [19.9 - 24.6] | 14.9 | [12.5 - 17.8] | 5.7 |
| **'CIP109994.scf.fasta'** | *Neisseria bacilliformis* ATCC BAA-1200 | 15.5 | [12.6 - 18.9] | 22.2 | [19.9 - 24.6] | 15.6 | [13.1 - 18.5] | 2.66 |
| **'CIP109994.scf.fasta'** | *Neisseria dumasiana* 93087 | 15.5 | [12.5 - 18.9] | 22.2 | [19.9 - 24.6] | 15.6 | [13.1 - 18.5] | 6.44 |

| Query | Subject | $d_0$ | C.I. $d_0$ | $d_4$ | C.I. $d_4$ | $d_6$ | C.I. $d_6$ | Diff. G+C Percent |
|---|---|---|---|---|---|---|---|---|
| **'CIP109994.scf.fasta'** | *Morococcus cerebrosus* CIP 81.93 | 14.8 | [11.9 - 18.2] | 22.2 | [19.9 - 24.7] | 15.0 | [12.5 - 17.9] | 5.51 |
| **'CIP103045.scf.fasta'** | *Neisseria macacae* ATCC 33926 | 14.7 | [11.8 - 18.1] | 22.1 | [19.9 - 24.6] | 14.9 | [12.4 - 17.7] | 5.73 |
| **'CIP109994.scf.fasta'** | *Eikenella exigua* PXX | 14.5 | [11.6 - 17.9] | 22.1 | [19.8 - 24.5] | 14.7 | [12.2 - 17.5] | 1.5 |
| **'CIP109994.scf.fasta'** | *Neisseria sicca* ATCC 29256 | 14.8 | [11.9 - 18.2] | 22.1 | [19.8 - 24.5] | 15.0 | [12.5 - 17.9] | 6.03 |
| **'CIP109994.scf.fasta'** | *Neisseria elongata* ATCC 25295 | 15.6 | [12.7 - 19.1] | 22.0 | [19.7 - 24.4] | 15.7 | [13.2 - 18.6] | 3.08 |
| **'CIP103045.scf.fasta'** | *Morococcus cerebrosus* CIP 81.93 | 14.7 | [11.8 - 18.1] | 22.0 | [19.7 - 24.4] | 14.9 | [12.4 - 17.7] | 5.54 |
| **'CIP103045.scf.fasta'** | *Neisseria dumasiana* 93087 | 15.8 | [12.8 - 19.3] | 21.9 | [19.6 - 24.3] | 15.9 | [13.3 - 18.8] | 6.48 |
| **'CIP103045.scf.fasta'** | *Neisseria sicca* ATCC 29256 | 14.7 | [11.9 - 18.1] | 21.9 | [19.6 - 24.3] | 14.9 | [12.5 - 17.8] | 6.06 |
| **'CIP109994.scf.fasta'** | *Uruburuella testudinis* DSM 26510 | 15.6 | [12.7 - 19.1] | 21.9 | [19.6 - 24.3] | 15.7 | [13.2 - 18.6] | 3.58 |
| **'CIP103045.scf.fasta'** | *Neisseria subflava* ATCC 49275 | 14.5 | [11.7 - 17.9] | 21.8 | [19.6 - 24.3] | 14.7 | [12.3 - 17.6] | 7.48 |
| **'CIP109994.scf.fasta'** | *Neisseria subflava* ATCC 49275 | 14.4 | [11.6 - 17.8] | 21.8 | [19.5 - 24.2] | 14.6 | [12.2 - 17.5] | 7.45 |
| **'CIP103045.scf.fasta'** | *Neisseria elongata* ATCC 25295 | 15.8 | [12.8 - 19.2] | 21.8 | [19.6 - 24.3] | 15.9 | [13.3 - 18.8] | 3.11 |
| **'CIP109994.scf.fasta'** | *Neisseria elongata* subsp. *glycolytica* ATCC 29315 | 15.7 | [12.8 - 19.2] | 21.7 | [19.4 - 24.1] | 15.8 | [13.3 - 18.7] | 2.91 |
| **'CIP103045.scf.fasta'** | *Uruburuella testudinis* DSM 26510 | 15.3 | [12.4 - 18.8] | 21.7 | [19.4 - 24.1] | 15.5 | [13.0 - 18.4] | 3.61 |
| **'CIP109994.scf.fasta'** | *Eikenella halliae* NML130454 | 14.5 | [11.7 - 17.9] | 21.7 | [19.5 - 24.2] | 14.8 | [12.3 - 17.6] | 0.65 |
| **'CIP103045.scf.fasta'** | *Neisseria elongata* subsp. *glycolytica* ATCC 29315 | 15.8 | [12.8 - 19.2] | 21.6 | [19.4 - 24.0] | 15.8 | [13.3 - 18.7] | 2.94 |
| **'CIP103045.scf.fasta'** | *Eikenella halliae* NML130454 | 14.5 | [11.6 - 17.8] | 21.5 | [19.3 - 23.9] | 14.7 | [12.2 - 17.5] | 0.68 |
| **'CIP103045.scf.fasta'** | *Eikenella corrodens* ATCC 23834 | 14.5 | [11.7 - 17.9] | 21.5 | [19.3 - 24.0] | 14.7 | [12.3 - 17.6] | 1.13 |
| **'CIP109994.scf.fasta'** | *Eikenella corrodens* ATCC 23834 | 14.5 | [11.7 - 17.9] | 21.5 | [19.2 - 23.9] | 14.7 | [12.3 - 17.6] | 1.1 |
| **'CIP103045.scf.fasta'** | *Uruburuella suis* DSM 17474 | 15.9 | [12.9 - 19.4] | 21.4 | [19.2 - 23.8] | 15.9 | [13.4 - 18.8] | 2.31 |
| **'CIP109994.scf.fasta'** | *Uruburuella suis* DSM 17474 | 15.8 | [12.8 - 19.2] | 21.4 | [19.2 - 23.8] | 15.8 | [13.3 - 18.7] | 2.27 |

#### Table 4: Strains in your dataset

Joint dataset of automatically determined closest type strains (if this mode was chosen), manually selected type strains (if selected accordingly) and the provided user strains, if provided (marked in yellow).

| Strain | Authority | Other deposits | Synonyms | Base pairs | Percent G+C | No. proteins | Goldstamp | Bioproject accession | Biosample accession | Assembly accession | IMG OID |
|---|---|---|---|---|---|---|---|---|---|---|---|
| *Neisseria weixii* DSM 103441 | Zhang et al. 2019 | 10022; CGMCC 1.15732 | *Neisseria weixii* | 2511 904 | 49.2 | 1927 | Gp0302531 | PRJNA394640 | SAMN07357044 | GCA_002327085 | |
| *Uruburuella testudinis* DSM 26510 | Kuhnert et al. 2015 | 07_OD624; CCUG 63373 | *Uruburuella testudinis* | 2949 123 | 53.3 | 2578 | | PRJNA224116 | SAMN24043414 | GCF_022870865 | |
| *Neisseria chenwenguii* 10023T | Zhang et al. 2019 | DSM 103440; CGMCC 1.15736 | *Neisseria chenwenguii* | 2496 444 | 54.0 | 2243 | Gp0261167 | PRJNA341560 | SAMN05726023 | GCA_002216145 | |
| *Eikenella exigua* PXX | Stormo et al. 2020 emend. Bernard et al. 2020 | DSM 109756; NCTC 14318 | *Eikenella exigua* | 1993 063 | 55.4 | 1960 | | PRJNA224116 | SAMN10995776 | GCF_008805035 | |
| *Neisseria subflava* ATCC 49275 | (Flügge 1886) Trevisan 1889 | LMG 5313; CCUG 23930; DSM 17610; CIP 103343; NRL 30,017; U37 | *Micrococcus subflavus*; *Neisseria subflava* | 2195 659 | 49.5 | 2063 | Gp0444348 | PRJNA224116 | SAMN11461056 | GCF_005221305 | |
| *Neisseria dumasiana* 93087 | Wroblewski et al. 2017 | LMG 30012; DSM 104677 | *Neisseria dumasiana* | 2639 953 | 50.5 | 2365 | | PRJNA224116 | SAMN06192246 | GCF_002108505 | |
| *Neisseria mucosa* ATCC 19696 | Véron et al. 1959 | CCUG 26877; DSM 17611; JCM 12992; NCTC 12978; CIP 59.51 | *Neisseria mucosa* | 2688 408 | 51.1 | 2386 | Gp0357104 | PRJNA445206 | SAMN08770273 | GCA_003028315 | |

| Strain | Authority | Other deposits | Synonyms | Base pairs | Percent G+C | No. proteins | Goldstamp | Bioproject accession | Biosample accession | Assembly accession | IMG OID |
|---|---|---|---|---|---|---|---|---|---|---|---|
| *Neisseria animalis* DSM 23392 | Berger 1960 | ATCC 49930; CCUG 808; NCTC 10212; CIP 72.15 | *Neisseria animalis* | 2174 610 | 51.9 | 2015 | | PRJNA394631 | SAMN07357035 | GCA_003795365 | |
| *Eikenella halliae* NML130454 | Bernard et al. 2020 | LMG 30894; NCTC 14180 | *Eikenella halliae* | 2538 675 | 56.3 | 2403 | | PRJNA310649 | SAMN04931641 | GCA_001648475 | |
| *Kingella potus* DSM 18304 | Lawson et al. 2005 | 3/SID/1128; CCUG 49773; NCTC 13336 | *Kingella potus* | 2327 523 | 57.9 | 2143 | Gp0013769 | PRJNA235010 | SAMN02746039 | | 2574180453 |
| *Uruburuella suis* DSM 17474 | Vela et al. 2005 | CCUG 47806; CECT 5685; strain 1258/02 | *Uruburuella suis* | 2635 278 | 54.7 | 2326 | Gp0325735 | PRJNA520322 | SAMN10866380 | GCA_004341385 | 2795385457 |
| *Neisseria dentiae* DSM 19151 | Sneath and Barrett 1997 | ATCC 700276; CIP 106968; Dent SHI/3848; V33 | *Neisseria dentiae* | 2758 789 | 54.4 | 2390 | | PRJNA224116 | SAMN06212708 | GCF_002108595 | |
| *Neisseria zoodegmatis* DSM 21643 | Vandamme et al. 2006 | LMG 23012; ATCC 29859; NCTC 12230; CDC D5986; CL 194/78 | *Neisseria zoodegmatis* | 2536 387 | 50.9 | 2296 | | PRJNA224116 | SAMN06212709 | GCF_002108575 | |
| *Neisseria elongata* subsp. *glycolytica* ATCC 29315 | Henriksen and Holten 1976 | CCUG 6508 A; CCUG 6508; DSM 23337; NCTC 11050; CIP 82.85 | *Neisseria elongata* subsp. *glycolytica* | 2275 949 | 54.0 | 3089 | Gp0003939 | PRJNA30471 | SAMN00008838 | GCA_000176755 | 647000283 |
| *Neisseria macacae* ATCC 33926 | Vedros et al. 1983 | DSM 19175; CIP 103346; M-740 | *Neisseria macacae* | 2682 989 | 51.2 | 3003 | Gp0006432 | PRJNA64729 | SAMN02299447 | GCA_000220865 | 651324075 |

| Strain | Authority | Other deposits | Synonyms | Base pairs | Percent G+C | No. proteins | Goldstamp | Bioproject accession | Biosample accession | Assembly accession | IMG OID |
|---|---|---|---|---|---|---|---|---|---|---|---|
| *Neisseria sicca* ATCC 29256 | (von Lingelsheim 1908) Bergey et al. 1923 | LMG 5290; CCUG 23929; CCUG 24959; DSM 17713; CIP 103345; NRL 30,016 | *Diplococcus siccus*; *Neisseria sicca* | 2830 772 | 50.9 | 3660 | Gp0003928 | PRJNA30481 | SAMN00008842 | GCA_000174655 | 645058748 |
| *Morococcus cerebrosus* CIP 81.93 | Long et al. 1981 | ACM 858; ATCC 33486; DSM 24335; NCTC 11393; UQM 858 | *Morococcus cerebrosus* | 2451 689 | 51.4 | 2795 | Gp0118031 | PRJNA267751 | SAMN03200160 | GCA_000813705 | |
| *Bergeriella denitrificans* NBRC 102155 | (Berger 1962) Xie and Yokota 2005 | ATCC 14686; CCUG 2155; DSM 17675; JCM 21446; NCTC 10295; CIP 72.16; IAM 14975 | *Bergeriella denitrificans*; *Neisseria denitrificans* | 2323 905 | 55.8 | 2118 | Gp0075760 | PRJDB1342 | SAMD00047216 | GCA_001592185 | |
| *Neisseria elongata* ATCC 25295 | Bøvre and Holten 1970 | LMG 5124; CCUG 2043; CCUG 2130 A; CCUG 2130; DSM 17712; NCTC 10660; CIP 72.27 | *Neisseria elongata*; *Neisseria elongata* subsp. *elongata* | 2359 626 | 53.8 | 2225 | Gp0127311 | PRJNA329891 | SAMN05421815 | GCA_900113545 | 2675903696 |
| *Eikenella corrodens* ATCC 23834 | (Eiken 1958) Jackson and Goodman 1972 | LMG 15557; CCUG 2138; DSM 8340; JCM 12952; NCTC 10596; CIP 70.75 | *Bacteroides corrodens*; *Eikenella corrodens* | 2129 788 | 55.8 | 2627 | Gp0003877 | PRJNA30493 | SAMN00008822 | GCA_000158615 | 643886208 |
| *Neisseria bacilliformis* ATCC BAA-1200 | Han et al. 2006 | CCUG 50858; DSM 23338; MDA 2833 | *Neisseria bacilliformis* | 2434 573 | 59.6 | 2966 | Gp0005240 | PRJNA53053 | SAMN00253995 | GCA_000194925 | 651324074 |

| Strain | Authority | Other deposits | Synonyms | Base pairs | Percent G+C | No. proteins | Goldstamp | Bioproject accession | Biosample accession | Assembly accession | IMG OID |
|---|---|---|---|---|---|---|---|---|---|---|---|
| CIP109994.scf.fasta | | | | 2183685 | 56.9 | 2032 | | | | | |
| CIP103045.scf.fasta | | | | 2291386 | 57.0 | 2110 | | | | | |

#### Methods, Results and References

The genome sequence data were uploaded to the Type (Strain) Genome Server (TYGS), a free bioinformatics platform available under https://tygs.dsmz.de, for a whole genome-based taxonomic analysis [1]. The analysis also made use of recently introduced methodological updates and features [2]. Information on nomenclature, synonymy and associated taxonomic literature was provided by TYGS's sister database, the List of Prokaryotic names with Standing in Nomenclature (LPSN, available at https://lpsn.dsmz.de) [2]. The results were provided by the TYGS on 2022-09-22. The TYGS analysis was subdivided into the following steps:

##### Determination of closely related type strains

Determination of closest type strain genomes was done in two complementary ways: First, all user genomes were compared against all type strain genomes available in the TYGS database via the MASH algorithm, a fast approximation of intergenomic relatedness [3], and, the ten type strains with the smallest MASH distances chosen per user genome. Second, an additional set of ten closely related type strains was determined via the 16S rDNA gene sequences. These were extracted from the user genomes using RNAmmer [4] and each sequence was subsequently BLASTed [5] against the 16S rDNA gene sequence of each of the currently 17188 type strains available in the TYGS database. This was used as a proxy to find the best 50 matching type strains (according to the bitscore) for each user genome and to subsequently calculate precise distances using the Genome BLAST Distance Phylogeny approach (GBDP) under the algorithm 'coverage' and distance formula $d_5$ [6]. These distances were finally used to determine the 10 closest type strain genomes for each of the user genomes.

##### Pairwise comparison of genome sequences

For the phylogenomic inference, all pairwise comparisons among the set of genomes were conducted using GBDP and accurate intergenomic distances inferred under the algorithm 'trimming' and distance formula $d_5$ [6]. 100 distance replicates were calculated each. Digital DDH values and confidence intervals were calculated using the recommended settings of the GGDC 3.0 [2,6].

##### Phylogenetic inference

The resulting intergenomic distances were used to infer a balanced minimum evolution tree with branch support via FASTME 2.1.6.1 including SPR postprocessing [7]. Branch support was inferred from 100 pseudo-bootstrap replicates each. The trees were rooted at the midpoint [8] and visualized with PhyD3 [9].

##### Type-based species and subspecies clustering

The type-based species clustering using a 70% dDDH radius around each of the 21 type strains was done as previously described [1]. The resulting groups are shown in Table 1 and 4. Subspecies clustering was done using a 79% dDDH threshold as previously introduced [10].

#### Results

##### Type-based species and subspecies clustering

The resulting species and subspecies clusters are listed in Table 4, whereas the taxonomic identification of the query strains is found in Table 1. Briefly, the clustering yielded 20 species clusters and the provided query strains were assigned to 1 of these. Moreover, user strains were located in 2 of 23 subspecies clusters.

##### Figure caption SSU tree

**Figure 1.** Tree inferred with FastME 2.1.6.1 [7] from GBDP distances calculated from 16S rDNA gene sequences. The branch lengths are scaled in terms of GBDP distance formula $d_5$. The numbers above branches are GBDP pseudo-bootstrap support values > 60 % from 100 replications, with an average branch support of 85.0 %. The tree was rooted at the midpoint [8].

##### Figure caption genome tree

**Figure 2.** Tree inferred with FastME 2.1.6.1 [7] from GBDP distances calculated from genome sequences. The branch lengths are scaled in terms of GBDP distance formula $d_5$. The numbers above branches are GBDP pseudo-bootstrap support values > 60 % from 100 replications, with an average branch support of 96.3 %. The tree was rooted at the midpoint [8].
